# PSK signaling controls ABA homeostasis and signaling genes and maintains shoot growth under osmotic stress

**DOI:** 10.1101/2020.10.20.347674

**Authors:** Komathy Rajamanickam, Martina D. Schönhof, Bettina Hause, Margret Sauter

**Author notes:** **Corresponding author**, Margret Sauter.

## Abstract

Water deficit impairs growth and survival of plants. Many water stress responses are under control of abscisic acid (ABA) but little is known about growth control under osmotic stress. Based on the previously described growth-promoting activity of the peptide hormone phytosulfokine (PSK), we hypothesized that it may contribute to growth regulation under water stress conditions. To test this hypothesis, we analyzed the *Arabidopsis thaliana* PSK receptor (PSKR) null mutant *pskr1-3 pskr2-1* under mannitol and drought stress. In particular under mild water stress, fresht weight and photosynthetic efficiency were more reduced in *pskr1-3 pskr2-1* than in wild type. Hydroponic and grafting experiments showed that PSKR signaling was not required for long-distance signaling from mannitol-stressed roots to shoot but rather for cell growth promotion in the shoot. Unlike wild type, *pskr1-3 pskr2-1* shoots did not accumulate ABA in response to mannitol, showed misregulation of ABA synthesis genes and elevated expression of *ABI1* and *ABI2*, repressors of ABA signaling whereas application of ABA partially reversed shoot growth inhibition by mannitol in *pskr1-3 pskr2-1*. In turn, mannitol and ABA induced expression of *PSK3* and PSKR1, and ABA promoted expression of *PSK2* and *PSK4* revealing feedback regulatory loops between PSKR and osmotic stress signaling.

**Highlight:** Phytosulfokine receptor signaling regulates ABA synthesis and signaling genes and promotes ABA accumulation in the shoot of water-stressed plants and maintains leaf growth and photosynthetic efficiency which ensures plant health.

## INTRODUCTION

Water deficit is a major challenge for plants as it impairs growth and ultimately survival. It is a result of osmotic stress brought about by drought or high salinity. The non-ionic osmolyte mannitol is frequently used in osmotic stress and plant growth research (Nikonorova *et al*., 2018; Kalve *et al*., 2020). Osmotic stress induces responses to ameliorate the stress conditions such as a reduction of water loss through stomatal closure (Munemasa *et al*., 2015). The plant hormone abscisic acid (ABA) accumulates in plant shoots following drought or osmotic stress and controls stoma closure and transcriptional changes (Bartels and Sunkar, 2005; Yamaguchi-Shinozaki and Shinozaki, 2006; Finkelstein, 2013, Takahashi *et al*., 2020). In addition, ABA-independent pathways of osmotic stress resistance exist.

Active ABA levels are determined by synthesis, degradation, inactivation and remobilization. ABA is synthesized from xanthophylls. Nine-*cis*-epoxycarotenoid dioxygenase (NCED) catalyze a key regulated step (Luchi *et al*., 2001; Schwartz *et al*., 2003) and aldehyde oxidases (AO) catalyze the conversion of abscisic aldehyde to ABA (Seo *et al*., 2000). Degradation of ABA is initiated by ABA 8’-hydroxylase, an enzyme that is encoded by four members of the cytochrome P450 CYP707A family in Arabidopsis (Kushiro *et al*., 2004; Saito *et al*., 2004). ABA hydroxylation has been identified as the key step in ABA catabolism (Dejonghe *et al*., 2018). However, inactivation of ABA also occurs through conjugation to glucose by ABA UDP-glucosyltransferase (UGT) encoded by *UGT71B6-B8*, in Arabidopsis (Dong *et al*., 2014). ABA-glucose ester is a storage form of ABA that can be remobilized by ABA glucosidases BG1 and BG2 (Lee *et al*., 2006). Thus, ABA homeostasis depends on synthesis and inactivation pathways whereby many of the genes involved are regulated in response to dehydration and other stresses (Xu *et al*., 2013).

Water deficit is first perceived in the root and from there communicated to the shoot. Among other drought-induced signals that move from root to shoot is the signaling peptide CLAVATA3/EMBRYO-SURROUNDING REGION-RELATED 25 (CLE25) that moves through the vasculature to leaves where it is perceived by BAM receptor-like kinases (Takahashi *et al*., 2018; Takahashi *et al*., 2019). CLE25/BAM signaling activates NCED3, a key gene in ABA biosynthesis and thereby contributes to ABA-dependent drought adaptation.The study revealed a crucial role of signaling peptides and receptor-like kinases in plant adaptation to water deficit.

Phytosulfokine (PSK) belongs to the group of secreted signaling peptides (Sauter, 2015; Kaufmann und Sauter, 2019). PSK is perceived by PSK receptors at the plasma membrane that belong to the leucine-rich repeat receptor-like kinase family encoded by two genes, PSKR1 and PSKR2, in *Arabidopsis thaliana* (Matsubayashi *et al*., 2006). Knockout of both receptor genes impairs root elongation and shoot growth whereas exposure of wild-type seedlings to PSK promotes growth by enhancing cell expansion (Kutschmar *et al*., 2009; Stührwohldt *et al*., 2011). At the molecular level, PSKR1 interacts with the co-receptor Brassinosteroid-insensitive (BAK1) and the H^+^-ATPases AHA1 and AHA2, and indirectly through BAK1 and AHAs with the Cyclic nucleotide-gated channel 17 (CNGC17). These proteins assemble in a nanocluster at the plasma membrane and were proposed to form a functional unit that drives cell expansion (Ladwig *et al*., 2015). PSKR signaling promotes protoplast expansion in a CNGC17-dependent manner and was proposed to lead to cell wall acidification, water uptake, accompanied by osmotic adjustment, and consequently to expansion of cells.

Low water potential prevents water uptake and limits cell expansion resulting in reduced growth rates under drought conditions. While physiological responses to water-deficit, foremost stoma closure, have been well studied, the question, if growth inhibition by osmotic constraint is balanced by a growth-promoting pathway to prevent an extreme stress response resulting in growth arrest, has not been resolved. It is clear however that an, albeit reduced, growth rate is maintained in plants despite of osmotic constraints. In support of the hypothesis that PSK signaling maintains plant growth under osmotic stress a recent study demonstrated that PSK precursor processing is required to promote root growth in response to the osmolyte mannitol in Arabidopsis (Stührwohldt *et al*., 2021).

Shoot growth is a more sensitive indicator to stress (Claeys *et al*., 2014) and was studied here to further investigate the role of PSKR signaling in response to osmotic stress. Our results show that PSKR signaling in the shoot is required to maintain shoot growth under mild osmotic stress, in part through altered ABA synthesis and signaling.

## MATERIALS AND METHODS

### Plant material and growth conditions

All experiments were performed with *Arabidopsis thaliana* (L.) Heynh. ecotype Columbia (Col-0) and mutants in the Col-0 background. The lines used were described previously as indicated: *pskr1-3* (Kutschmar *et al*., 2009; Stührwohldt *et al*., 2011), *pskr2-1* (Amano *et al*., 2007; Stührwohldt *et al*., 2011), *pskr1-2pskr2-1* (Stührwohldt *et al*., 2011; Hartmann *et al*., 2013), *PSKR1ox2* and *PSKR1ox12* (Hartmann *et al*., 2013), *35S:PSKR1-GFP* (Hartmann *et al*., 2015). For GUS analyses of PSK receptors the lines *PSKR1:GUS-4* (Kutschmar *et al*., 2009; Stührwohldt *et al*., 2011) and *PSKR2:GUS-3* were used. For growth on plates, seeds were surface-sterilized for 25 min with 2% (w/v) sodium hypochlorite (NaOCl) followed by four washing steps with autoclaved water and placed on square plates containing half-strength MS medium (Murashige & Skoog, 1962; basal salt mixture, Duchefa Biochemie) and 1% (w/v) sucrose, solidified with 0.4% Gelrite (Duchefa Biochemie). After two days of stratification at 4°C in the dark, plates were transferred to long day conditions with a 16 h light (70 μM photons m^−2^ s^−1^) and 8 h dark cycle at 22°C and 60% humidity.

For germination and greening assays, wild type and mutant parental plants were grown and harvested at the same time to ensure equal seed quality. Seeds were placed on medium supplemented with or without 1 μM *(+)-cis, trans-abscisic* acid (ABA) (Duchefa Biochemie), 350 μM mannitol and/or 1 μM PSK (Pepscan). PSK and ABA were always added freshly. ABA was diluted right before use. The penetration of the endosperm or testa by the embryo radicle was counted as successful germination event. Three independent experiments were performed with 100 seeds each per genotype per treatment.

To analyze shoot growth under stress conditions, seedlings were pregrown vertically under sterile conditions for 4 days to complete germination and cotyledon greening (Supplemental Fig. S6) and subsequently transferred to new plates supplemented with mannitol, sorbitol and/or ABA as indicated and grown for the times indicated. Growth was quantified with a fine scale as shoot fresh weight or shoot dry weight. Plant pictures are shown on a black background for better visualization. Primary root lengths were determined 7 days after transfer of 4-day-old seedlings to mannitol because the roots reached the bottom of the plate after longer times making it impossible to measure root lengths lateron.

For grafting, 6-day-old seedlings were grown on agar plates (0.5X MS pH 5.8; 0.5% (w/v) sucrose; 1% (w/v) agar) at short-day conditions (8 h light and 16 h dark). Grafting was performed as described (Marsch-Martínez *et al*., 2013). The rootstock and scion were prepared under sterile conditions by excising the hypocotyl of the seedlings. Cotyledons were also excised to improve healing. Adventitious roots were removed immediately when they appeared. The grafted plants were allowed to recover for 14 days and successful grafts were chosen for osmotic stress experiments with mannitol. Plants were grown on manitol or mannitol-free medium as indicated was for another 14 days prior to analysis.

### Drought experiment and plant analysis

Plants were grown in square pots (7×7×8 cm) on soil for three weeks under well-watered conditions with 50 ml of water every third day. After three weeks, the soil was water-saturated for 3 days and, subsequently, plants were watered every 3 days with 5 ml for another 3 weeks. Pots with control plants remained well-watered as before. Drought experiments were repeated three times and growth was quantified as shoot fresh weight.

For chlorophyll fluorescence measurements, an IMAGING-PAM chlorophyll fluorometer (Maxi version with blue measuring light, Walz, Effeltrich, Germany) was used. Plants were dark-adapted for 30 minutes prior to image capture. The maximum quantum yield of photosystem II (PS II) was measured as the ratio of *Fv/Fm* = (*Fm* – *Fo*)/*Fm*, where *Fo* is the minimum fluorescence measured using a weak excitation beam (setting 1 with 1 Hz), Fm is the maximum fluorescence measured by applying a saturated light pulse (2,500 μmol photons m^−2^ s^−1^) and Fv is the variable fluorescence (Fm-Fo). Subsequently, plants were exposed to saturating pulses with background illumination and ΔF/Fm’ measurement was done under steady state conditions. ΔF/Fm’ was calculated using the formula (Fm ‘-F)/Fm’, where the prime (‘) denotes actinic light. Images and measurements were obtained using Imaging Win V2.41a software (Walz).

### Histochemical GUS analysis

To analyze spatial distribution of *PSKR* and *PSK* expression *β*-glucuronidase (GUS) assays were performed as described (Weigel & Glazebrook, 2002) with minor changes. Seedlings were collected in 90% (v/v) isopropanol, incubated for 10 minutes and washed with 50 mM sodium phosphate buffer (pH 7.2). Staining was performed for 15 h and stopped by transferring the seedlings to 70% (v/v) ethanol. Tissues were cleared with a chloral hydrate:deionized water:glycerol 6:2:1 (g/ml/ml) mixture that was added on microscopy slides instead of water. Seedlings were visualized under bright-field illumination with a Nikon SMZ18 binocular (Nikon) and photographed with a DIGITAL SIGHT-Ri1 camera (Olympus).

### RNA isolation, RT-PCR and quantitative real time PCR

Total RNA was isolated from true leaves using TRI Reagent (Merck) following manufacturer’s protocol. cDNA was synthesized from 1 μg DNase I-treated (Thermo Fisher Scientific) total RNA by oligo(dT)-primed reverse transcription using RevertAid Reverse Transcriptase (Thermo Fisher Scientific). To avoid gDNA contamination and to ensure an equal amount of RNA input, cDNA was tested by PCR for *ACTIN2* (At3g18780) transcripts prior to qPCR analysis with the primers ACT2for 5’-CAAAGACCAGCTCTTCCATCG-3’ and ACT2rev 5’-CTGTGAACGATTCCTGGACCT 3’. qPCR was used to examine the expression of *PSKR1, PSKR2, PSK1, PSK2, PSK3, PSK4, PSK5, NCED3, NCED5, NCED9, CYP707A1, CYP707A2, CYP77A3, CYP707A4, UGT71B6, UGT71B7, UGT71B8, BG1, BG2, AAO1, AAO2, ABI1* and *ABI2* with primers listed in Supplemental Table 1 using the Rotor Gene SYBR Green PCR Kit (Qiagen) according to manufacturers’ protocol in a Rotor gene Q cycler (Qiagen). Ten ng cDNA per sample were applied in a total volume of 15 μl. Estimation of raw data was done with the Rotor-Gene Q 2.3.1.49 (Qiagen) program.

The relative transcript abundance was calculated based on the ΔΔCP method including primer efficiency to normalize the data with two reference genes (Pfaffl, 2001; Van Desompele *et al*., 2007). In relation to each reference gene (*ACT2* and GLYCERALDEHYDE-3-PHOSPHATE DEHYDROGENASE 1 (*GAPC1*)) (Supplemental Table 1) values were averaged from three independent biological replicates with two technical replicates each. The expression of wild type under control conditions was set to 1 and all other values were calculated as fold change of that.

### ABA measurement

Four-day-old seedlings were transferred to medium containing mannitol as indicated and grown for additional 7 days. Roots and shoots were harvested separately and immediately frozen in liquid nitrogen. About 50 mg of homogenized, frozen material was extracted with 500 μl of methanol containing isotope-labelled internal standard ^2^H_6_-ABA (0.1 ng μl^−1^) followed by centrifugation. The supernatant was diluted with 4.5 ml water and subjected to solid-phase extraction on HR-XC (Chromabond, Macherey-Nagel, Düren, Germany) column. Fractions containing ABA were eluted with acetonitrile and separated using the ACQUITY UPLC System (Waters, Eschborn, Germany) (Balcke *et al*. 2012). Detection of ABA and ^2^H_6_-ABA was done by ESI-tandem mass spectrometry (MS/MS) using a 3200 Q TRAP^®^ LC/MS/MS mass spectrometer (Waters) (Balcke *et al*., 2012). ABA content per sample was calculated using the ratio of ABA and ^2^H_6_-ABA peak heights. Data were obtained from 5 biological replicates each.

To an alternative method to measure changes in ABA levels in response to mannitol, we obtained the *ABAleon2.1* line and the plasmid *barII-UT-ABAleon2.1* from Rainer Waadt (Waadt *et al*., 2014) that we used to generate a homozygous *pskr1-3pskr2-1 ABAleon2.1* line. Plants were transformed with *Agrobacterium tumefaciens EHA105*, containing the *barII-UT-ABAleon2.1* construct with the floral dip method (Clough and Bent, 1998). The progeny were selected with BASTA^®^. In the T2 generation the seedlings were additionally screened with the FastGene^®^ Blue/Green LED Flashlight (Nippon Genetics Europe GmbH) with an orange/blue filter to exclude silencing of the transgene. Acceptor photobleaching Förster resonance energy transfer (FRET) was used to measure relative differences in ABA levels in each genotype using a Leica SP5 CLSM. Cells were excited sequentially at 458 and 514 nm and emission recorded with adequate filter sets. Post-bleach images were captured at 458 nm excitation (Supplemental Fig. S8). FRET is visualized as an increase in mTurquoise fluorescence following cpVenus173 photobleaching (Supplemental Figure S8). FRET efficiency was calculated according to the formula FRET_eff_ = (D_post_-D_pre_)/D_post_ with D_post_ =fluorescence intensity of the donor after acceptor photobleaching and D_pre_ =fluorescence intensity of the donor before acceptor photobleaching.

### Statistical analysis

Statistical analyses were done using Minitab^®^ 16.1. Data that were not distributed normally were evaluated using a Kruskal-Wallis or Mann-Whitney test for pairwise comparison. Normally distributed data were tested for equal variance. In the case of equal variance, ANOVA, otherwise a t-test for pairwise comparison was chosen.

### Accession numbers

*ACT2* - At3g18780, *GAPC1* - At3g04120, *PSKR1* - At2g02220, *PSKR2* - At5g53890, *PSK1* - At1g13590, *PSK2* - At2g22860, *PSK3* - At3g44735, *PSK4* - At3g49780, *PSK5* - At5g65870, *NCED3* - At3g14440, *NCED5* - At1g30100, *NCED9* - At1g78390, *CYP707A1* - At4g19230, *CYP707A2* - At2g29090, *CYP707A3* - At5g45340, *CYP707A4* - At3g19270, *UGT71B6* - At3g21780, *UGT71B7* - At3g21790, *UGT71B8* - At3g21800, *BG1* - At1 g52400, *BG2* - At2g32860, *AAO1* - At5g20960, *AAO2* - At3g43600, *ABI1* - At4g26080, *ABI2* - At5g57050.

## RESULTS

### PSKR1 promotes shoot growth under osmotic stress

PSK signaling is known to promote root and shoot growth (Sauter, 2015). A recent report demonstrated that PSK precursor synthesis and precursor processing by subtilisin serine proteases enhance root growth under mannitol stress conditions revealing a role of PSK signaling of growth under osmotic stress (Stührwohldt *et al*., 2021). To better understand PSK signaling of growth under osmotic stress, we employed the PSK receptor null line *pskr1-3 pskr2-1* (Stührwohldt *et al*., 2011; Hartmann *et al*., 2013) and focussed on shoot growth of seedlings exposed to mannitol. Seedlings were grown for 4 days without mannitol, then transferred to plates containing 0, 25 mM, 50 mM, 100 mM or 200 mM mannitol. Shoot growth was analyzed after an additional 2 weeks (Fig. 1). Wild type seedlings displayed inhibition of shoot size, shoot fresh weight and shoot dry weight by mannnitol in a dose-dependent manner, a phenotype that was exacerbated in *pskr1-3 pskr2-1* seedlings (Fig. 1A, B, C and D). The strongest growth inhibition of *pskr1-3 pskr2-1* seedlings compared to wild type was observed at low concentrations of 25 mM and 50 mM mannitol suggesting that PSKR signaling maintains shoot growth particularly well during mild osmotic stress. Growth reduction induced by mannitol was largely due to reduced leaf size (Fig. 1B). To see if cell expansion was dependent on PSK signaling under mannitol stress as reported for unstressed conditions (Matsubayashi *et al*., 2006; Kutschmar *et al*., 2009; Stührwohldt *et al*., 2011; Ladwig *et al*., 2015) we analyzed epidermal cell sizes of wild type and *pskr1-3 pskr2-1* first true leaves (Fig. 1E and F). The average epidermal cell size was smaller in *pskr1-3 pskr2-1* compared to wild type at control conditions (Fig. 1E, F). When exposed to 50 mM mannitol, cells became even smaller in both genotypes with a more severe effect observed in *pskr1-3 pskr2-1* seedlings indicating that PSK receptor signaling promotes cell expansion under osmotic stress conditions but cannot fully overcome the stress. As of note, reduction in cell size went along with reduced lobing of the epidermal cells under mannitol (Fig. 1E). The osmotic compound sorbitol that was used for comparison also exacerbated growth inhibition in *pskr1-3 pskr2-1* seedlings compared to wild type in a dose-dependent manner (Supplemental Fig. S1). Primary root growth was inhibited at 200 mM mannitol in both genotypes with a significantly stronger inhibition in *pskr1-3 pskr2-1* than wild type confirming previously reported results (Stührwohldt *et al*., 2021) whereas root elongation was unaffected by mannitol up to 100 mM in both genotypes (Supplemental Fig. S2) indicating that PSKR signaling was important to maintain shoot at mild osmotic stress conditions.

**Figure 1.**
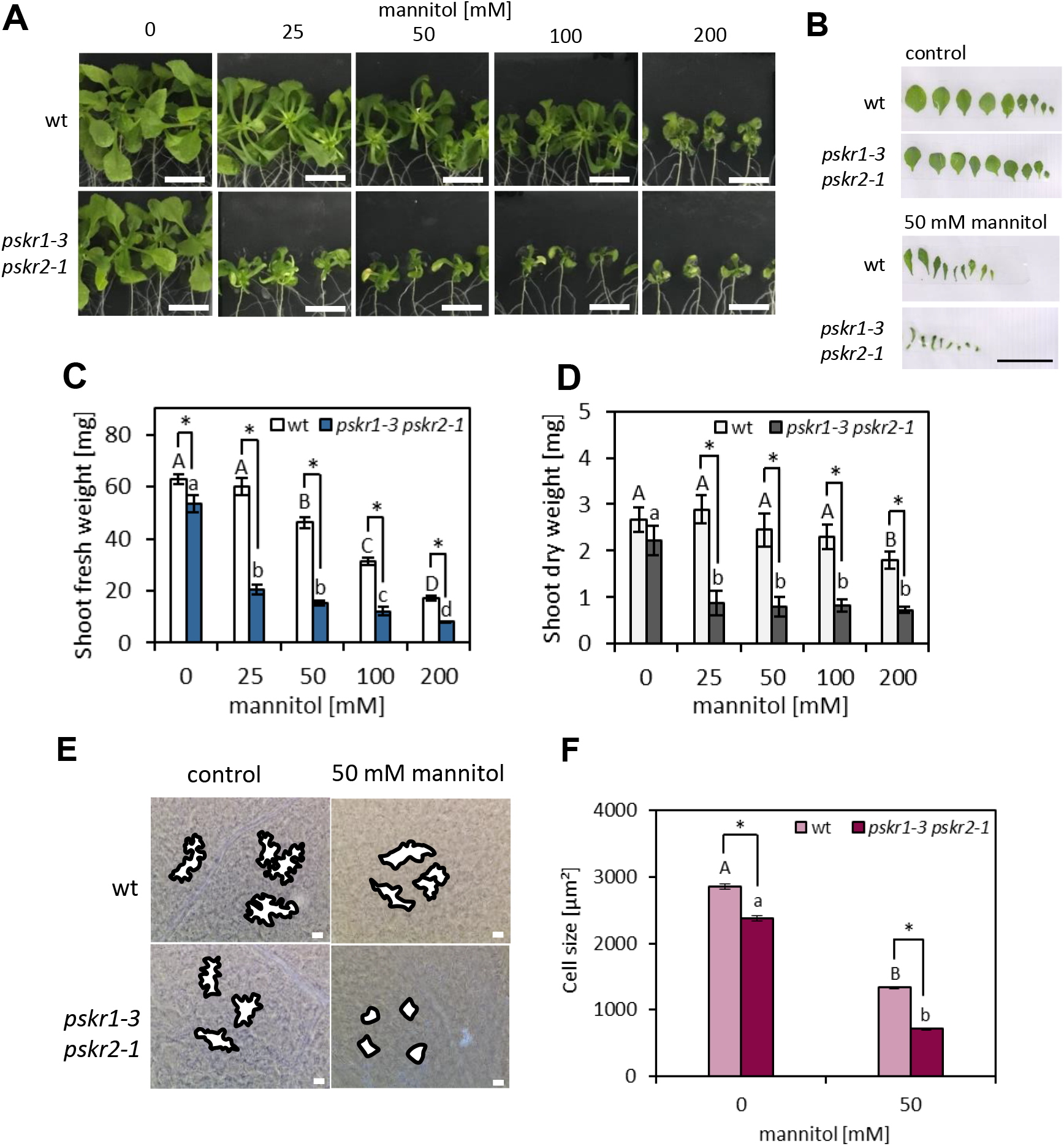
PSK receptor signaling promotes shoot growth under mannitol stress. Four-day-old wild type and *pskr1-3 pskr2-1* seedlings were transferred to media containing mannitol as indicated. (A) Shoot phenotypes of wild type (wt) and *pskr1-3 pskr2-1* plants after 2 weeks growth on mannitol. Scale bars = 10 mm. (B) Leaf numbers, sizes and morphology of wild type and *pskr1-3 pskr2-1* plants grown on mannitol-free medium or on 50 mM mannitol for 2 weeks. Scale bar = 10 mm. (C) Average shoot fresh weights (±SE) of plants grown as in A. (D) Shoot dry weights of plants grown as in A. Significantly different values within a genotype are denoted by different letters (Kruskal-Wallis, Tukey’s test, *p*<0.05, n≥36, 3 independent experiments). Asterisks indicate significant differences between genotypes (Mann-Whitney test, *p*<0.05). (E, F) Size of epidermal cells at the abaxial side. Plants were grown as in A-D. Single cell outlines are highlighed in bold black lines (scale bar = 10 nm). Average cell sizes (±SE) of the first true leaf were analyzed from 12 seedlings with 40-50 cells measured per seedling. Significant differences between treatments are indicated by different letters. Asterisks indicate significant differences between genotypes (Mann-Whitney test, *P*<0.05, n≥1500, 3 independent experiments).

To pinpoint which of the two PSK receptors mediated resistance to osmotic stress-induced growth inhibition of the shoot, we used single PSK receptor gene knock out lines (Kutschmar *et al*., 2009). Growth inhibition by 50 mM mannitol was comparable in the single knockout line *pskr1-3* and the double knockout line *pskr1-3 pskr2-1* whereas *pskr2-1* plants showed wild-type shoot growth indicating that PSKR1 promotes growth of plant shoots exposed to mild osmotic stress (Fig. 2).

**Figure 2.**
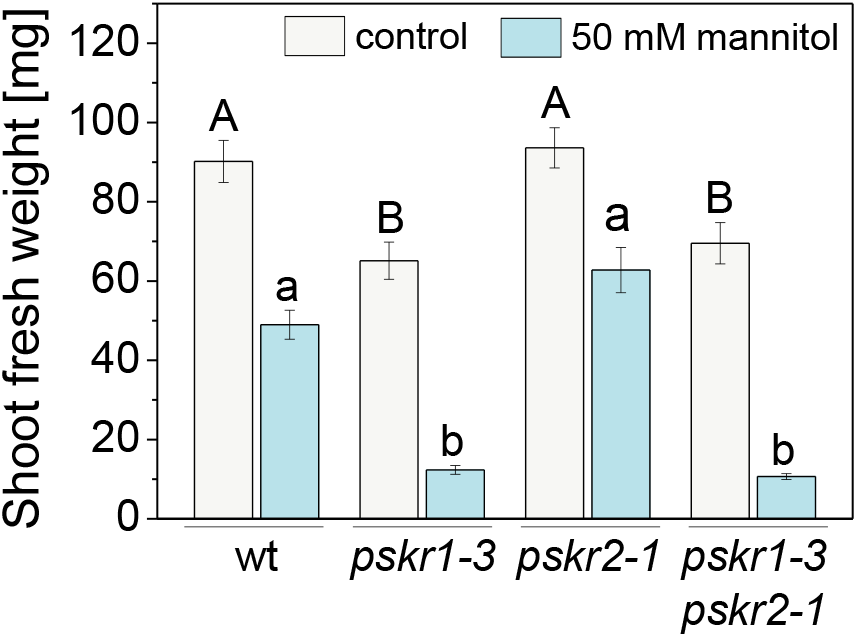
*PSKR1* is crucial for growth promotion under mannitol stress. Four-day-old wild type, *pskr1-3, pskr2-1*, and *pskr1-3 pskr2-1* seedlings were transferred to media with or without 50 mM mannitol. Mean (±SE) shoot fresh weights of seedlings after 3 weeks. Different letters indicate significant differences (Kruskal-Wallis, Tukey’s test, *p*<0.05, n≥36, 3 independent experiments).

For shoot growth experiments, we grew seedlings for 4 days on medium without osmoticum prior to osmotic stress treatment. However, osmotic stress resistance may also be important at earlier developmental stages during germination. To close this knowledge gap, we investigated seed germination and cotyledon greening in wild type and *pskr1-3 pskr2-1* exposed to mannitol, PSK and ABA. ABA is known as a drought hormone and mediates osmotic stress responses (Zhao *et al*., 2018). While seed germination was delayed by ABA and mannitol, no significant differences in germination rate were observed between wild type and *pskr1-3 pskr2-1* at any of the treatments indicating that inhibition of seed germination by ABA or mannitol was not controlled by PSK/PSKR signaling (Supplemental Figs. S3 and S4). On the other hand, cotyledon greening that is known to be delayed under unfavourable conditions such as under ABA or mannitol (Guan *et al*., 2014) was delayed in wild type but not *pskr1-3pskr2-1* seedlings (Supplemental Figs. S5 and S6) suggesting that PSK/PSKR signaling contributes to stress acclimation during very early seedling development. Application of mannitol to four-day-old seedlings was hence a useful approach to circumvent these early developmental processes when studying shoot growth regulation.

### PSKR signaling in the root is not required for osmotic stress signaling to the shoot

Water deficit is perceived in roots and the stress signal is transmitted to the shoot via the vasculature (Takahashi and Shinozaki, 2019; Takahashi *et al*., 2020). We next clarified whether PSKR signaling was required for stress perception in the root or for a root-derived stress signal in the shoot. Osmotic stress was routinely applied by transfering seedlings to plates containing mannitol. In order to test whether PSKR signaling of growth was induced through direct contact of leaves with mannitol or as part of the water deficit response signaled by roots, we compared growth on plates where the whole seedling had access to media to growth on plates where only roots were in contact with media (Fig. 3A, B). Stronger inhibition of shoot growth was observed in *pskr1-3 pskr2-1* compared to wild type in both setups but shoots that were not in contact with media began to dry out after eight days (Fig. 3A and B). To overcome this problem, a hydroponic system was established (Fig. 3C) where seedlings were grown on a mesh separating the shoot from the medium. After 10 days, seedlings were transferred to media containing 50 mM mannitol and grown for another 14 days. Shoot growth of *pskr1-3 pskr2-1* seedlings was significantly impaired compared to wild type (Fig. 3D-F) indicating that osmotic stress sensed by the roots led to PSKR-dependent shoot growth promotion.

**Figure 3.**
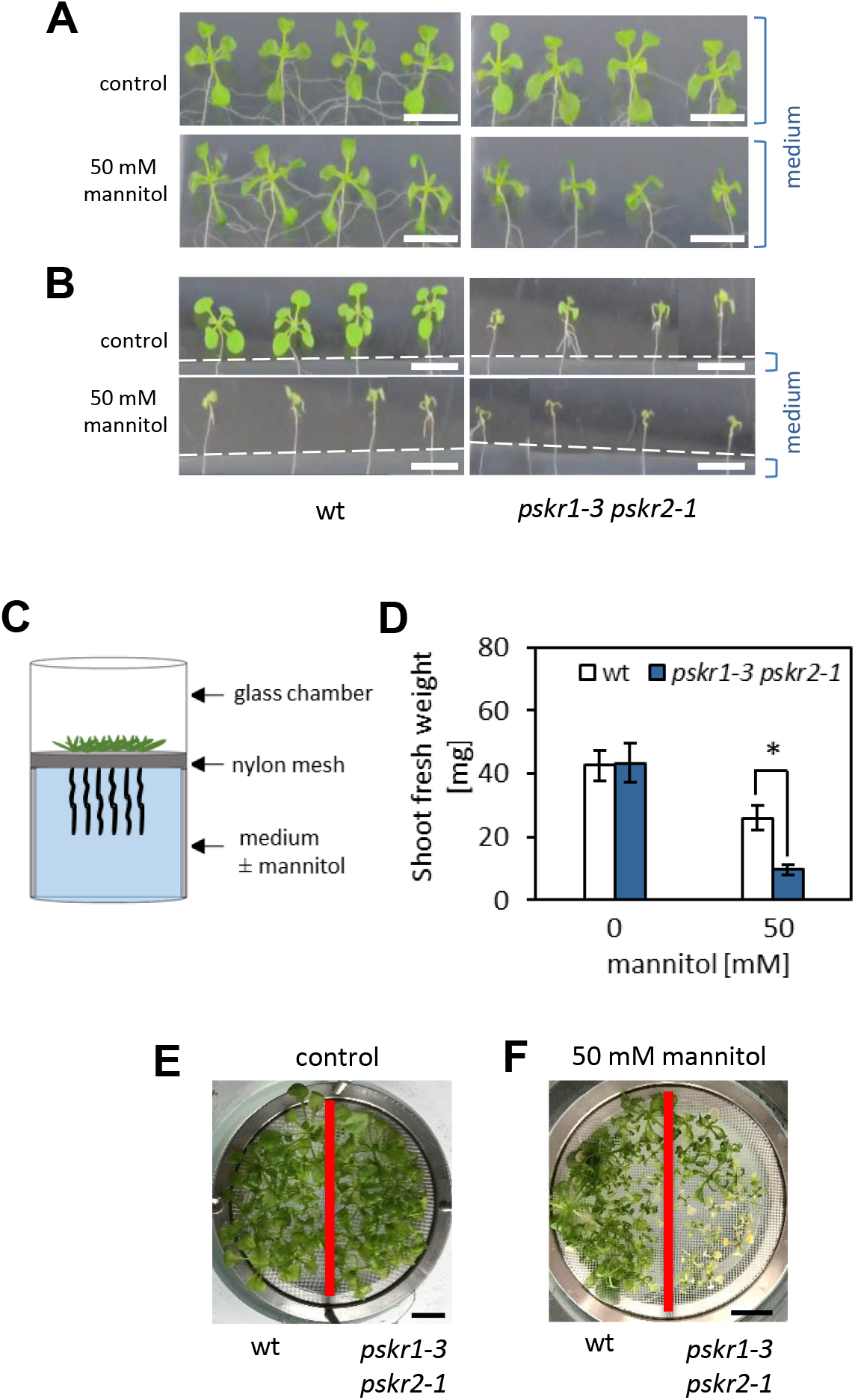
Mannitol sensing in roots is independent of *PSKRs*. (A) Seedlings were pregrown for 4 days and then transferred to medium supplemented with or without 50 mM mannitol. Phenotypes of shoots grown on plates with media for 8 days. (B) Seedlings were grown on plates with excised media at the height of the shoot to prevent direct contact of the shoot with mannitol (A,B: scale bar = 10 mm). (C) Schematic of the hydroponic system used to apply mannitol to roots. (D) Seedlings were pre-grown for 10 days as shown in C and then transferred to fresh medium supplemented with or without 50 mM mannitol. Average shoot fresh weights (±SE) were determined after 2 weeks from 3 independent experiments (Mann-Whitney test, *P*<0.05, n≤45). (E, F) Shoot phenotypes of plants grown on medium (E) without or (F) with 50 mM mannitol (scale bar = 10 mm).

To find out if PSKR signaling was required for osmotic stress perception in roots and participated to signal water deficit to the shoot, we performed grafting experiments as a widely used technique to investigate long-distance signaling in plants (Corbesier *et al*., 2007; Chen *et al*., 2006; Molnar *et al*., 2010; Liang *et al*., 2012). Wild type and *pskr1-3 pskr2-1* shoot scions and root stocks were grafted as indicated schematically in Figure 4A. Cotyledons were removed before grafting to prevent formation of adventitious roots which lower grafting efficiency. Ungrafted seedlings, also with cotyledons removed for better comparison, and within-genotype self-grafts were included as controls. Grafted wild type and *pskr1-3 pskr2-1* seedlings showed the same growth phenotypes as ungrafted seedlings indicating that the setup worked properly (Fig. 4C-E). Seedlings with a wild type shoot and a *pskr1-3 pskr2-1* root had the same shoot growth phenotype as wild type seedlings at control and stress conditions whereas seedlings with a *pskr1-3 pskr2-1* shoot and a wild type root had a *pskr1-3 pskr2-1* phenotype (Fig. 4C-E). These observations indicated that PSKR signaling does not contribute to root-shoot communication of osmotic stress but is required for growth promotion in the shoot during the stress. Excision of cotyledons inhibited shoot growth of *pskr1-3 pskr2-1* but not wild type seedlings even in the absence of osmotic stress (compare Fig. 1A, C and Fig. 4A, E) suggestive of a crosstalk between cotyledons and true leaves that is dependent on PSKR signaling. Taken together, the results showed that PSKR acts downstream of a mobile root-derived signal in the shoot to maintain growth under osmotic stress.

**Figure 4.**
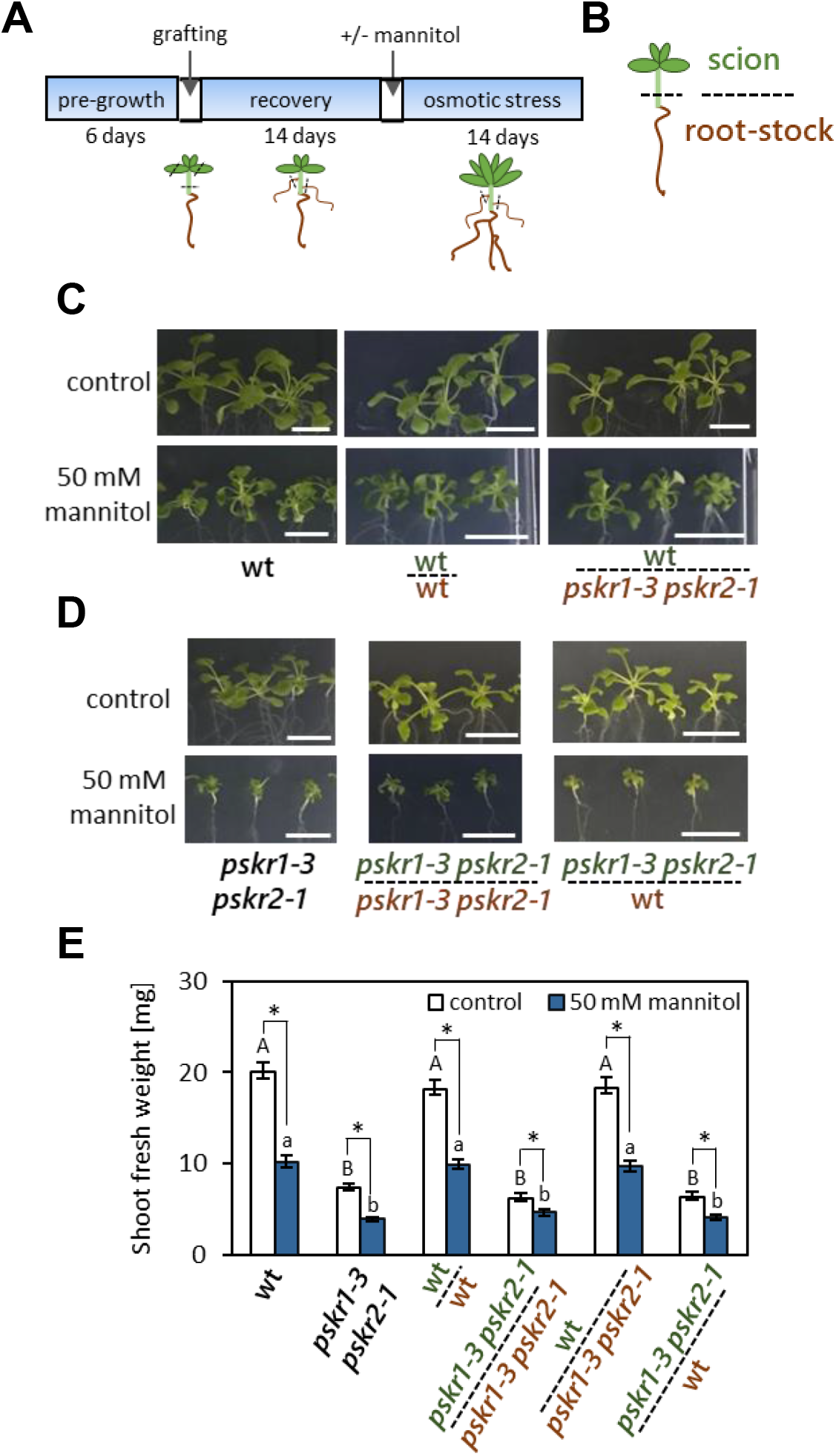
PSKR signalling in the shoot promotes growth in response to mannitol. Wild type shoots were grafted with double receptor knockout roots (wt + *pskr1-3 pskr2-1*) and vice versa (*pskr1-3 pskr2-1* + wt) under short day conditions. Successful grafts were analyzed for shoot fresh weight grown under long day conditions. (A) scheme representing the grafting procedure. (B) Model to represent labelling. (C, D) Shoot phenotypes of successful grafts on (±) 50 mM mannitol plates. Cotyledons of wt and *pskr1-3 pskr2-1* were cut and included as controls (scale bar = 10mm). (E) Average shoot fresh weights (±SE) of plants supplemented with/without 50 mM mannitol were determined after 2 weeks. Results are averages obtained from three independent experiments. The capital and minor letters indicate significant differences between genotypes upon control and mannitol treatments respectively (Kruskal-Wallis test *P*<0.05, n≥24) and asterisk indicate significant differences within treatments for a particular genotype (Mann-Whitney test, *P*<0.05)

To confirm that the PSKR-dependent maintenance of shoot growth under mild mannitol stress was a response to low water potential, we exposed wild type and *pskr1-3 pskr2-1* seedlings to mild drought stress (Fig. 5A). Plants were grown for 3 weeks under well-watered conditions and subsequently watered with one tenth the water supplied before. Shoot fresh weight after 3 weeks of growth under water-limiting conditions was reduced by about 50% in *pskr1-3 pskr2-1* seedlings compared to wild type (Fig. 5B). Enhanced drought symptoms in the knock-out mutant were further revealed by reduced photosynthetic efficiency of PSII (Fig. 5C, D). Using IMAGING-PAM, two parameters in dark-adapted and steady-state light conditions, Fv/Fm and ΔF/Fm’, were measured that reveal a response to abiotic stress (Zhang and Sharkey 2009; Kalaji *et al*., 2016), including drought (Yao *et al*., 2018). From the false colour images (Fig. 5C) the average Fv/Fm and ΔF/Fm’ ratios of rosettes were determined in both genotypes under control and drought conditions. The Fv/Fm ratio was close to 0.8 in both genotypes in well-watered conditions and decreased more under drought in *pskr1-3 pskr2-1* than wild type (Fig. 5D). The ΔF/Fm’ ration indicates steady-state quantum yield of PSII in light-adapted conditions and is an accurate indicator of operational PSII efficiency (Murchie and Lawson 2013). This ratio was also significantly lower in drought-exposed *pskr1-3 pskr2-1* plants compared to wild type (Fig. 5E) revealing that plants lacking PSKR signaling are more sensitive to low water potential.

**Figure 5:**
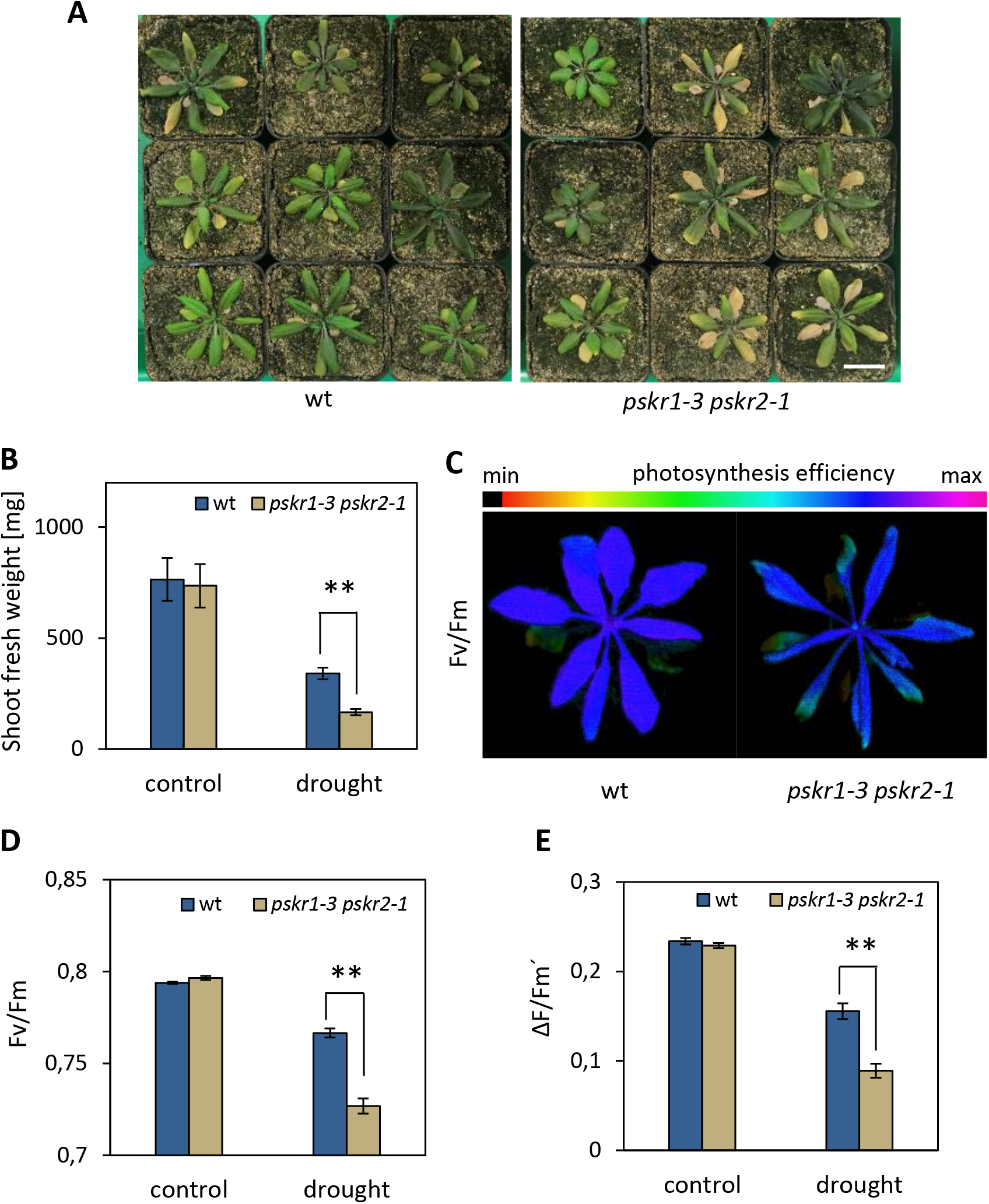
PSK receptor signaling provides drought stress tolerance. (A) Phenotypes of plants exposed to mild drought stress (scale bar = 3 cm). Plants were watered with 50 ml every 3 days for 3 weeks, soil water-saturated for 3 days and subsequently watered with 5 ml every 3 days for another three weeks or with 50 ml as a control. (B) Average (±SE) shoot fresh weights of wild type and *pskr1-3 pskr2-1* plants obtained from three independent experiments (Mann-Whitney test, *P*<0.01, n≥36). (C) Representative chlorophyll fluorescence images of drought exposed plants displaying the maximum quantum yield of PS II, Fv/Fm. (D) Fv/Fm ratio (±SE) of control and drought-stressed wild type and mutant plants (Mann-Whitney test, \*\**P*<0.001, n≥36). (E) ΔF/Fm’ ratio (Mann-Whitney test, \*\**P*<0.01, n≥36).

### Osmotic stress and ABA promote expression of *PSKR* and *PSK* genes

We next analyzed expression of *PSK* and *PSKR* genes in response to mannitol and ABA as a drought stress-induced hormone. *Promoter:GUS* lines (Kutschmar *et al*., 2009; Stührwohldt *et al*., 2011) were used to obtain spatial resolution and microarray and qRT-PCR data were used to quantify changes in gene expression. *PSKR1:GUS* activity was detected in cotyledons, true leaves and in the root and appeared to be induced in true leaves by mannitol (Fig. 6A-F). Microarray data showed induction of *PSKR1* in shoots after 1 day of treatment with 300 mM mannitol (Fig. 6H). RT-qPCR data confirmed elevated *PSKR1* transcript levels in true leaves after 1 day of exposure to 50 mM or 200 mM mannitol as well as by 1 μM and 5 μM ABA (Fig. 6I). By contrast, *PSKR2:GUS* activity was present in the cotyledon hydathode region (Fig. 6G) and *PSKR2* was induced in true leaves at 200 mM mannitol and 5 μM ABA but not at lower concentrations (Fig. 6J) in accord with the finding that shoot growth promotion under mild osmotic stress was dependent on *PSKR1* (Fig. 2).

**Figure 6.**
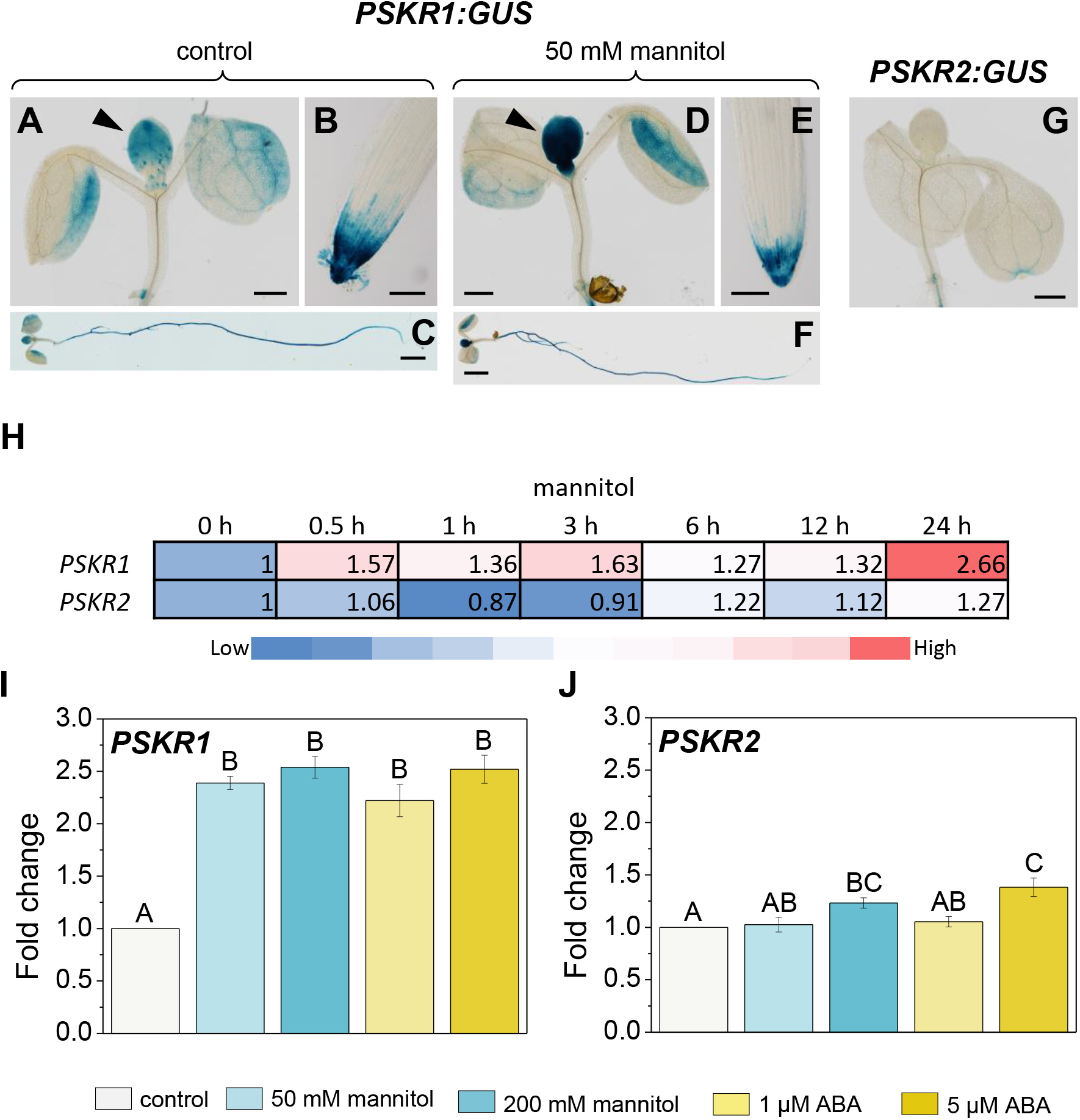
*PSKR1* expression is induced by mannitol and ABA. (A-F) *PSKR1:GUS-4* activity in 7-day-old seedlings treated with 50 mM mannitol for 1 day or left untreated as a control. (A-C) Expression in shoot, and root under control conditions. (D-F) Expression in shoot, and root under mannitol. Arrowheads point at true leaves; scale bar = 0.5 mm (A, D), = 0.1 mm (B, E), = 2 mm (C, F). (G) *PSKR2:GUS-3* activity in the shoot; scale bar = 0.5 mm. (H) Shoot tissue samples of 18-day-old plants analyzed for the expression of *PSKR1* and *PSKR2* under 300 mM mannitol treatment. Microarray data obtained from Arabidopsis efp browser 2.0 database available online (http://www.bar.utoronto.ca/) (Kilian *et al*., 2007). (I, J) Seven-day-old seedlings exposed to 50 mM, 200 mM mannitol, 1 μM or 5 μM ABA for 1 day or left untreated as a control. Relative transcript levels of (H) *PSKR1* and (I) *PSKR2* were analyzed by RT-qPCR in true leaves. Values are means (±SE); different letters indicate significant differences (one-way ANOVA, Tukey’s test, *P*<0.05, 3 biological replicates).

Of the five PSK precursor genes, *PSK2, PSK3, PSK4* and *PSK5* but not *PSK1* were expressed in the shoot in accord with the previous finding that *PSK1* is root-specific (Kutschmar *et al*., 2009) (Fig. 7A). *PSK3:GUS* activity was particularly high in young, expanding leaves. Analysis of PSK precursor gene expression by RT-qPCR showed significant induction of *PSK3* by 200 mM mannitol and 5 μM ABA in wild type while *PSK2* and *PSK4* expression was induced by ABA (Fig. 7B-F). Knockout of PSK signaling in *pskr1-3 pskr2-1* resulted in elevated *PSK2* and *PSK4* expression at control conditions suggestive of feedback inhibition by PSKR signaling. Mannitol but not ABA resulted in hyperinduced *PSK1, PSK2, PSK3* and *PSK5* transcript levels in *pskr1-3 pskr2-1* compared to wild type. Taken together, the data revealed common and differential induction of PSK genes in response to mannitol and ABA and a negative feedback loop between PSKR signaling and *PSK* expression.

**Figure 7.**
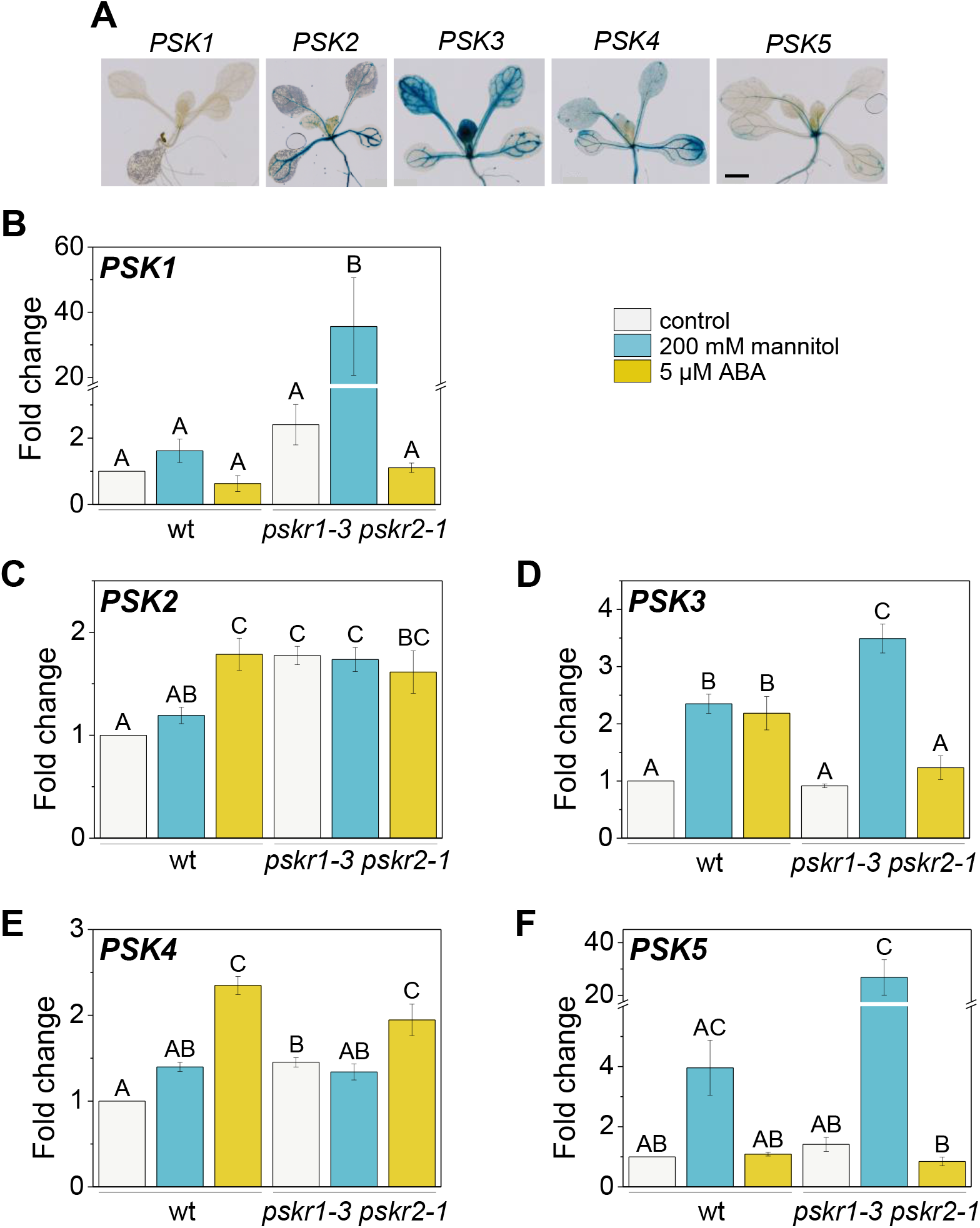
*PSK* genes are differentially regulated by mannitol, ABA and *PSKR* signaling. (A) GUS staining of *PSK1:GUS-3, PSK2:GUS-2, PSK3:GUS-3, PSK4:GUS-4* and *PSK5:GUS-1* seedlings; scale bar = 2 mm. (B-F) Seven-day-old seedlings were exposed to 200 mM mannitol or 5 μM ABA for 1 day for qPCR analyses in first true leaves. Results (±SE) are averages from 3 biological replicates with two technical repeats each. Different letters indicate significant differences. (B) Transcript levels of PSK genes relative to *PSK3* that was set to 1. (C-F) Relative transcript levels of *PSK1-5* in wild type and in *pskr1-3 pskr2-1* seedlings treated with mannitol or ABA as indicated (B, F: Kruskal-Wallis, Tukey’s test, *P*<0.05; C-E: one-way ANOVA, Tukey’s test, *P*<0.05).

### PSKR signaling regulates ABA levels and ABA responsiveness

Responses to osmotic stress are regulated by ABA-dependent and ABA-independent pathways. *PSK3* and *PSKR1* were regulated by both, mannitol and ABA suggesting that PSK/PSKR-ABA crosstalk occurs under osmotic stress. Since ABA is predominantly synthesized in leaves upon drought (Cardoso *et al*., 2020) we asked whether ABA levels were regulated by PSKR signaling. Analysis of ABA levels in wild type seedling shoots by mass spectrometry revealed a dose-dependent increase in ABA in response to mannitol (Fig. 8A). A similar increase was observed in roots (Supplemental Fig. S7). By contrast, ABA levels did not increase in *pskr1-3 pskr2-1* shoots in response to mannitol whereas in roots, ABA levels increased in *pskr1-3 pskr2-1* seedlings at 200 mM mannitol albeit overall levels were lower than in wild type (Supplemental Fig. S7). To verify a PSKR-dependent ABA accumulation in the shoot, we introduced the ABA sensor ABAleon2.1 (Waadt *et al*. 2014) into the *pskr1-3 pskr2-1* background. Photobleaching-based FRET analysis (Supplemental Fig. S8) revealed a decreased FRET efficiency, reporting elevated ABA in wild type but not in *pskr1-3 pskr2-1* in response to mannitol (Fig. 8B). The difference in absolute FRET efficiency between genotypes may be owed to the fact that the lines were transformed rather than introgressed.

**Figure 8.**
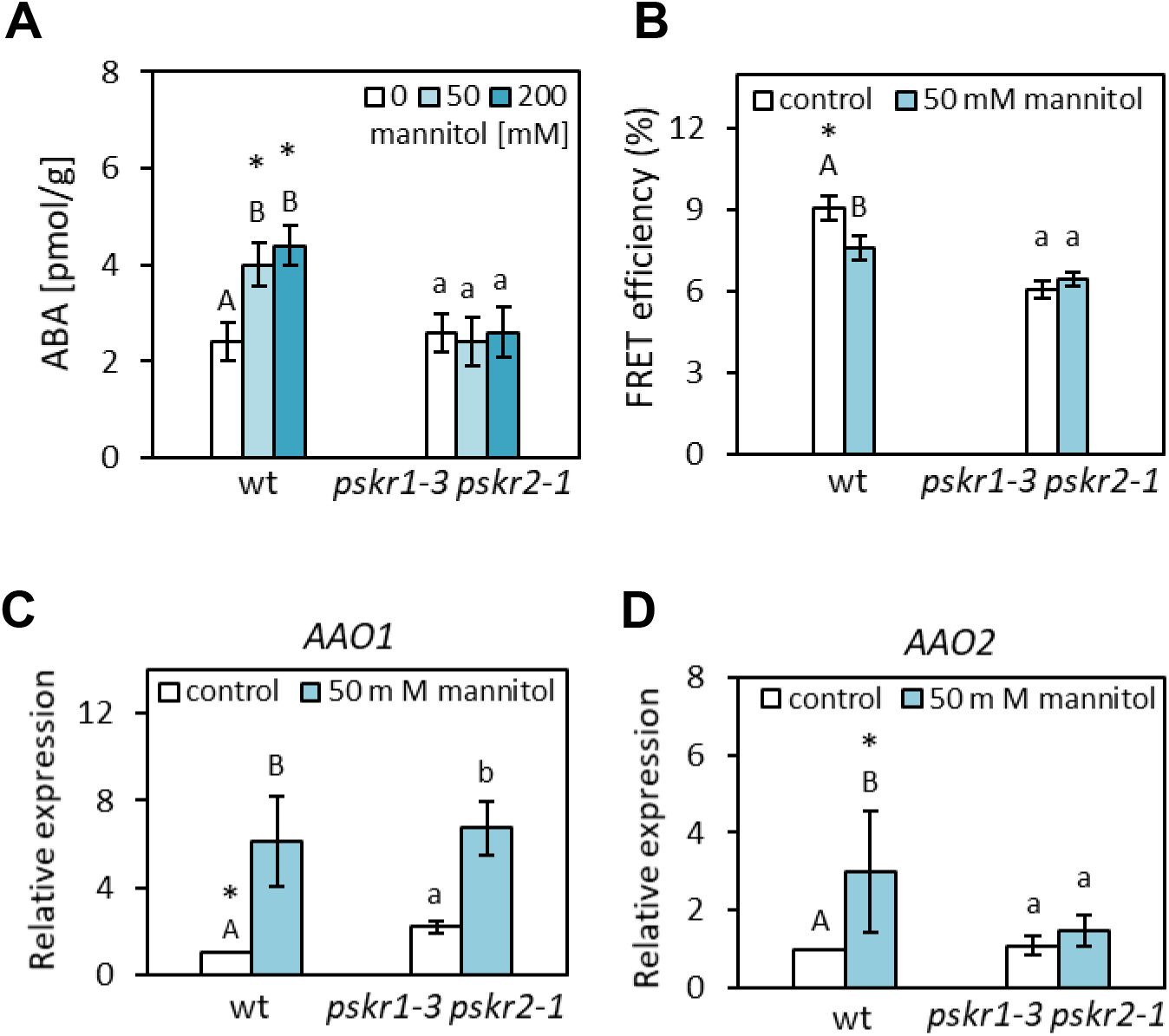
Mannitol-induced ABA accumulation in the shoot is PSKR-dependent. (A) Four-day-old wild type and *pskr1-3 pskr2-1* seedlings were transferred to 0 mM, 50 mM or 200 mM mannitol and ABA levels in the shoot were determined after 7 days. Different capital and minor letters indicate statistically significant differences between treatments (Kruskal-Wallis, Tukey’s test, *P*<0.05, 5 biological replicates). Asterisks indicate significant differences between genotypes per treatment (Mann-Whitney test, *P*<0.05, n=6). (B) ABA levels in the first true leaf were compared by photobleaching-based FRET analysis in wild type and *pskr1-3 pskr2-1* seedlings treated as in A. Decreased FRET efficiency indicates elevated ABA levels. Values are averages (±SE). Different capital and minor letters indicate significantly different values between treatments and asterisk indicate significant differences between genotypes for a specific treatment (Mann-Whitney test, *P*<0.05, n?142, 3 independent experiments). (C, D) Transcript of the ABA synthesis genes *AAO1* and *AAO2* were determined by RT-qPCR in true leaves of 4-day-old seedlings transferred to mannitol for another 3 days as indicated. Significantly different values for wt and *pskr1-3 pskr2-1* are represented by capital (one-way ANOVA with Tukey’s test, *P*<0.05, n=6) and minor letters (Mann-Whitney test, *P*<0.05, n=6). Asterisks indicate significant differences between genotypes at given treatment (Mann-Whitney test, *P*<0.05, n=6).

NCED3 is an ABA-biosynthetic enzyme that is upregulated in water-stressed leaves (Iuchi *et al*., 2001; Endo *et al*., 2008). We analyzed expression of *NCED3, NCED5* and *NCED9*, but found no significant differences in shoots of wild type and *pskr1-3 pskr2-1* seedlings exposed to mannitol (Supplemental Fig. S9). We next analyzed *AAO* gene expression. AAO enzymes catalyze the conversion of abscisic aldehyde to ABA. *AAO1* transcript levels were significantly higher in *pskr1-3 pskr2-1* than wild type and were upregulated in both genotypes by mannitol to the same level (Fig. 8C). By contrast, *AAO2* was upregulated significantly by mannitol in wild type but not *pskr1-3 pskr2-1* (Fig. 8D) which may contribute to the lower ABA level observed under osmotic stress.

Apart from synthesis, ABA levels depend on transient in-/activation and on degradation. ABA is inactivated to ABA-glucosyl ester (ABA-GE) by glycosyl transferases UGT71B6-B8 (Priest *et al*., 2006) (Supplemental Figure S9A). Of these, UGT71B6 is induced by mannitol in wild type and hyperinduced by mannitol in *pskr1-3 pskr2-1*, suggesting that loss of PSKR signaling favors ABA inactivation under mannitol stress. No genotype-dependent differences were observed in response to ABA. Release of ABA from ABA-GE is catalyzed by two ABA glucosidases, BG1 and BG2 (Xu *et al*., 2012). Transcripts of *BG1* were induced by ABA but not mannitol whereas *BG2* was not regulated. No differences in response to mannitol or ABA were observed between wild type and *pskr1-3 pskr2-1* (Supplemental Figure S10). *CYP707A1-4* genes code for ABA 8’-hydroxylases that degrade ABA to phaseic acid (Kushiro *et al*., 2004; Saito *et al*., 2004; Okamoto *et al*., 2006). *CYP707A1* transcript levels were higher in *pskr1-3pskr2-1* than in wild type exposed to mannitol possibly indicating enhanced ABA degradation in the mutant under osmotic stress (Supplemental Figure S11). In conclusion, the findings revealed crosstalk between PSKR signaling and ABA metabolism and suggest that PSKR signaling promotes expression of genes that favor ABA accumulation. These observations are in accord with the finding that PSKR signaling is needed to enhance ABA levels in response to mannitol.

We next explored whether the inability of *pskr1-3 pskr2-1* seedling shoots to accumulate ABA under osmotic stress could be complemented by exogenous ABA. A dose-response analysis showed that ABA at 0.1 μM and higher inhibited shoot growth in wild type under control conditions whereas *pskr1-3 pskr2-1* seedling growth was inhibited at 0.3 μM ABA and higher (Fig. 9A, B, C and D). In the presence of mannitol, shoot growth was promoted at 0.01 μM ABA in wild type and at 0.1 and 1 μM ABA in *pskr1-3 pskr2-1* seedlings indicating that low levels of ABA promote shoot growth under osmotic stress. The need for higher ABA required to promote shoot growth in *pskr1-3 pskr2-1* seedlings may be explained by a lower endogenous ABA content fostering the idea that PSKR promotes growth in part by increasing ABA in response to mannitol. It should be noted, however, that growth inhibition by mannitol was only partially alleviated by ABA in *pskr1-3 pskr2-1* suggesting that other levels of regulation exist.

**Figure 9.**
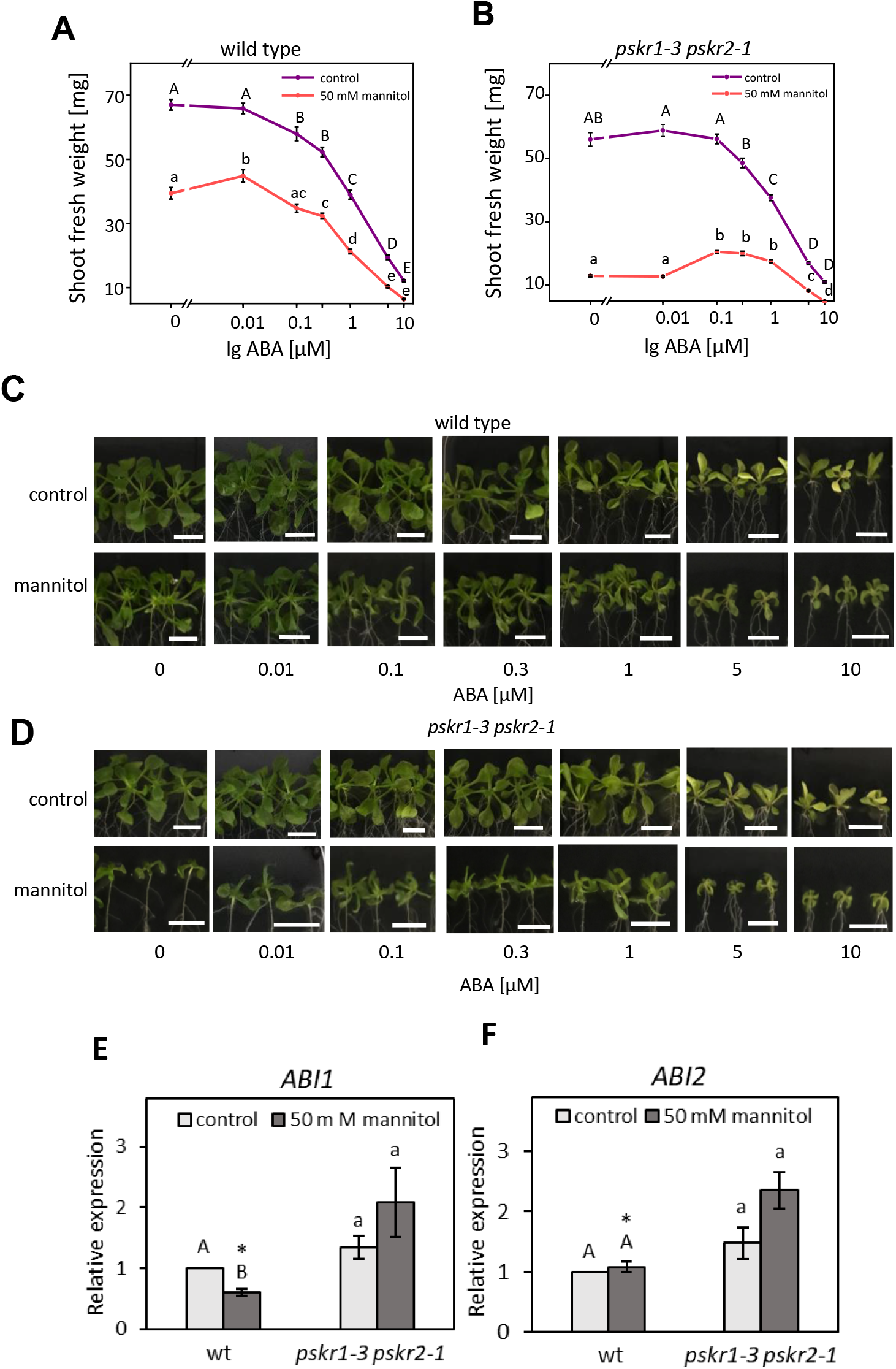
PSKR and ABA pathways partially interact. (A, B) Shoot fresh weights of wild type and *pskr1-3 pskr2-1* seedlings that were transferred after 4 days to plates containing ABA and mannitol as indicated were determined after 2 weeks (Kruskal-Wallis, Tukey’s test, *P*<0.05; n=36). (C, D) Phenotypes of seedlings analyzed in A, B. (E, F) Relative expression (±SE) of *ABI1* and *ABI2* in true leaves of 4-day-old seedlings grown on mannitol for another 3 days. Significantly different values for wild type and *pskr1-3 pskr2-1* are represented by capital and minor letters (Mann-Whitney test, *P*<0.05, n=6). Asterisks indicate significant differences between genotypes at a given treatment (Mann-Whitney test, *P*<0.05, n=6).

Finally, we analyzed a possible interaction between PSKR signaling and ABA signaling by using two marker genes of ABA signaling, *ABA-INSENSITIVE 1 (ABI1)* and *ABI2*, that act as negative regulators of ABA signaling and have been implicated in stress acclimation (Ludwików, 2015). Under mannitol, *pskr1-3 pskr2-1* seedlings accumulated higher *ABI1* and *ABI2* transcript levels than wild type whereas no difference was observed at unstressed conditions (Fig. 9E, F) indicating that ABA signaling under osmotic stress may be repressed in *pskr1-3 pskr2-1*. In conclusion, several lines of evidence suggest a role for PSKR signaling in plant acclimation to osmotic stress through crosstalk with ABA metabolism and signaling at the transcriptional level (Figure 10).

**Figure 10.**
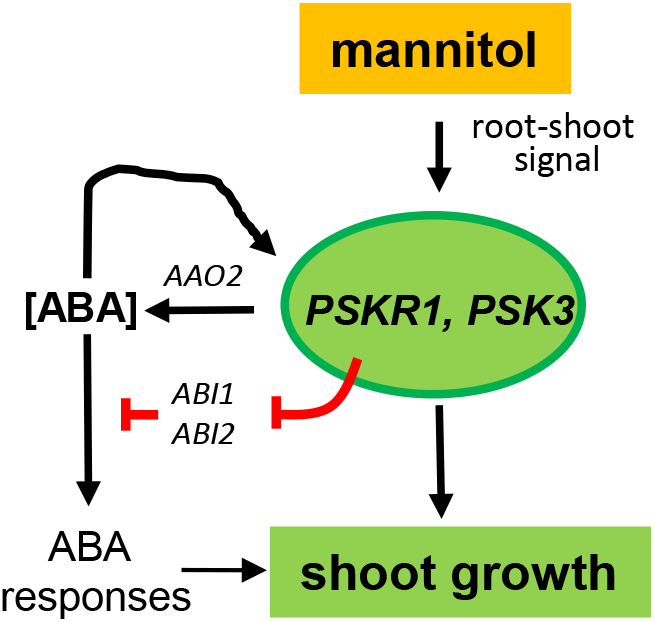
Model of shoot growth control during mannitol stress by PSKR1 and ABA. Mannitol induces *PSKR1* and *PSK3* gene expression and PSKR signaling promotes ABA synthesis, through upregulation of the ABA synthesis gene *AAO2*, and ABA signaling, through repression of *ABI1* and *ABI2*. In mutants that lack PSKRs, reduced ABA in mannitol-stressed shoots contributes to shoot growth inhibition. Growth under mannitol stress is further promoted by ABA-independent PSKR signaling.

## DISCUSSION

### PSK receptor signaling mediates osmotic stress-induced inhibition of cotyledon development

Seed germination and seedling establishment are stringently controlled by environmental conditions to ensure seedling survival. In particular, water availability is a requirement for seedling establishment. Under water-limited conditions, ABA acts as a hormonal signal that prevents germination and inhibits seedling growth. PSK signaling was previously shown to promote organ growth through cell expansion (Kutschmar *et al*., 2009; Hartmann *et al*., 2013). Under osmotic stress imposed by mannitol treatment, PSK receptor signaling inhibited postgermination seedling development while the PSK receptor null mutant *pskr1-3 pskr2-1* showed enhanced cotyledon greening and seedling growth. Seedling development is likewise inhibited by ABA. Both, mannitol and ABA-dependent inhibition were, to a large part, dependent on PSKR signaling. More so, seedlings overexpressing PSKR1 were hypersensitive to ABA and to mannitol suggesting that PSKRs mediate growth adaptation in response to osmotic stress. Inhibition of cotyledon greeening by PSKR signaling is a novel finding as PSKRs were previously described as growth-promoting receptors. The current observation thus extends our view on growth regulation by PSKRs and reveals that PSKR signaling inhibits early seedling growth under unfavorable environmental conditions. By contrast, postgermination shoot growth under water-limiting conditions was promoted by *PSKR1*. PSKR-dependent inhibition of growth at the early seedling stage and promotion of postgermination shoot growth under water-limiting conditions suggests that developmental arrest of seedlings and maintenance of vegetative growth of plants are best choices to cope with water shortage and that PSKR signaling can be wired accordingly.

### PSKRs support shoot growth during osmotic stress

A recent report showed that PSK precursor processing via SBT3.8 subtilase improves root growth and drought stress resistance in Arabidopsis (Stührwohldt *et al*., 2021). Drought stress experiments described here revealed that *PSKR* signaling is required to maintain postgermination shoot growth, and photosynthetic efficiency to counteract drought-induced senescence. It was previously reported that *PSKR1* delays senescence after bolting and provides cellular longevity and potential for growth (Matsubayashi *et al*., 2006). This ability appears particularly important during water limiting conditions. A crucial role of *PSKR1* in promoting osmotic stress resistance of the shoot was revealed in single receptor gene knockout lines. *pskr1-3* seedlings displayed the same growth retardation as seedlings of the double receptor knock out line *pskr1-3 pskr2-1* whereas *pskr2-1* seedlings had a wild type phenotype.

Grafting experiments and osmotic stress application to the root showed that shoot growth requires PSKR signaling in the leaves likely induced by a long-distance stress signal transmitted from the roots via the vasculature (Takahashi and Shinozaki, 2019; Takahashi *et al*., 2020). Whether the peptide PSK participates in long distance signaling cannot be excluded but, clearly, PSKR signaling is neither required for signal synthesis in the root nor for the transmission of the signal from root to shoot. In the leaves, cells communicate via the extracellular signal PSK and the plasma membrane-bound PSKR1 receptor despite of a continuous network of plasmodesmata that enable cell-cell communication. It has been suggested that apoplastic signals allow for signal integration and output coordination (Chivasa and Goodman, 2020). In that light, PSK/PSKR1 signaling appears highly suitable to coordinate leaf growth.

Plants adjust shoot and root growth to limited soil water availability with different sensitivities dependent on the degree and duration of the stress (Deak and Malamy, 2005; Comas *et al*., 2013; Pierik and Testerink, 2014; Koevoets *et al*., 2016). Root growth was inhibited at 200 mM mannitol and higher (this study; Stührwohldt *et al*., 2021) whereas shoot growth was reduced already at 25 mM mannitol supporting the finding that shoot growth is more sensitive to the stress (Claeys *et al*., 2014). In accord with a well-adjusted stress management, shoot growth maintenance via PSKR1 signaling was most efficient at low mannitol concentrations providing particularly good protection against mild water stress to the highly sensitive shoot.

### PSK precursor and receptor genes are induced in the shoot by mannitol and ABA to promote growth

Plant shoots exposed to mannitol displayed elevated transcript levels of *PSKR1* and of the PSK precursor gene *PSK3*, that were also induced by ABA suggesting that the PSK signal pathway is promoted by osmotic stress signaling through ABA to actively maintain shoot growth. Mannitol and ABA exerted common but also differential control of PSK signaling genes pointing to a fine-tuned regulatory network rather than a linear pathway of growth control. *PSK2* and *PSK4* were induced by ABA but not by mannitol revealing ABA regulation of PSK signaling in a pathway that is unrelated to mannitol-induced osmotic stress. Intricate regulatory loops were revealed by hyperinduction of *PSK1, PSK3* and *PSK5* by mannitol in the PSK receptor null background compared to wild type which suggested that PSK signaling feedback-inhibits expression of ligand precursor genes under osmotic stress possibly to balance growth under water-limited conditions. The regulated signaling genes *PSK3* and *PSKR1* were expressed most prominently in young expanding leaves in accord with their role in driving cell expansion (Hartmann *et al*., 2013; Ladwig *et al*., 2015).

### ABA homeostasis in osmotically stressed leaves is controlled by PSKR signaling

Mannitol-induced osmotic stress led to an increase of ABA in leaves. The elevation of ABA levels was dependent on PSKR signaling raising the question how PSK signaling controls ABA homeostasis at the molecular level. The steady-state level of active ABA is determined by ABA synthesis, catabolism, inactivation and remobilization (Cutler and Krochko, 1999; Nambara and Marion-Poll, 2005). NCEDs are key enzymes of ABA synthesis that cleave 9-*cis*-xanthophylls to xanthoxin, a precursor of ABA. Arabidopsis NCED3 was previously described as the major stress-induced *NCED* gene in leaves with *NCED5* and *NCED9* being induced to a minor degree in response to water-deficit (Tan *et al*., 2003). In our study, *NCED3* was induced by mannitol in wild type and hyperinduced in the *PSKR* null mutant whereas *NCED5* was induced by ABA which favors elevated ABA synthesis in drought-stressed shoots in both genotypes. The final step in ABA synthesis is catalyzed by AAO that is encoded by two genes in Arabidopsis, *AAO1* and *AO2* (Seo *et al*., 2000). *AAO1* was induced by mannitol in wild type and *pskr1-3 pskr2-1* shoots, whereas *AAO2* was induced in wild type but not *pskr1-3 pskr2-1* indicating that at least some of the capacity to synthesize ABA in response to osmotic stress is controlled by *PSKRs*. Similarly, *pskr1-3pskr2-1* shoots showed hyperinduction of the ABA-conjugating gene *UGT71B6* and downregulation of the ABA-glucose β-glucosidase gene *BG1* in shoots when exposed to mannitol. BGs were previously described as key regulators of osmotic stress resistance (Xu *et al*., 2012). *BG1* is regulated by miR165/166 and downregulation of miR165/166 resulted in elevated *BG1* expression and in elevated ABA levels (Yan *et al*., 2016). Taken together, the data show that PSKR signaling impacts expression of genes related to ABA metabolism in response to mannitol. The changes in gene expression suggest that ABA synthesis and release from ABA-glucose may be reduced in the PSK receptor null mutant whereas conjugation of ABA may be favored. This would explain why ABA levels do not increase in osmotically stressed *pskr1-3 pskr2-1* shoots.

In addition to ABA synthesis, ABA signaling may be controlled by *PSKRs*. ABA is perceived by Pyrabactin Resistant/Pyrabactin Resistant-Like/Regulatory Components of ABA Receptor (PYR/PYL/RCAR) receptors. The protein phosphatases 2C (PP2Cs) ABI1 and ABI2 act as PYR/PYL/RCAR-coreceptors that keep downstream SnRK2s in an inactive state in the absence of ABA (Joshi-Saha *et al*., 2011; Mitula *et al*., 2015). When PYR/PYL/RCAR receptors bind ABA, the PP2Cs are inactivated leading to the activation of SnRKs that are critical for osmotic stress responses (Boudsocq *et al*., 2004; Fujii *et al*., 2011). Expression analysis revealed higher transcript levels of the negative ABA regulator genes *ABI1* and *ABI2* in *pskr1-3 pskr2-1* compared to wild type following mannitol treatment suggesting that *PSKRs* promotes not only synthesis but also ABA signaling during osmotic stress.

A particular role of ABA in growth maintenance under water stress conditions was revealed by exogenous ABA. In wild type, ABA promoted shoot growth at 0.01 μM whereas the growth-promoting concentrations were shifted to 0.1-1 μM ABA in *pskr1-3 pskr2-1* seedlings. A requirement for a higher concentration of ABA ineed to promote growth is in accord with lower endogenous ABA levels in the mutant. The relative growth promoting effect of ABA under mannitol stress was higher in the mutant than in wild type but ABA did not fully restore the dwarf phenotype of the mutant at any concentration indicating that additional pathways are misregulated in *pskr1-3 pskr2-1* seedlings that ensure growth under water-limiting conditions.

In summary, PSKR signaling controls adaptation of the shoot to drought stress by maintaining growth under water-limiting conditions and by maintaining photosynthetic activity. PSKR signaling is not required for long-distance signaling of osmotic stress from the root to the shoot but rather promotes ABA levels and possibly ABA signaling in the shoot. Our study reveals a role of PSKR signaling in balancing shoot growth and water-stress responses.

## SUPPLEMENTARY DATA

Table S1. Primer sequences used for qRT-PCR.

Fig. S1. PSKRs confer enhanced resistance to sorbitol.

Fig. S2. PSK receptor signaling is required to maintain root growth under severe osmotic stress.

Fig. S3. Inhibition of seed germination by mannitol is not dependent on PSK receptor signaling.

Fig. S4. Inhibition of seed germination by ABA is not dependent on PSK receptor signaling.

Fig. S5. Repression of early seedling development by ABA or mannitol is mediated by PSK receptor signaling.

Fig. S6. Inhibition of cotyledon greening by ABA is partially dependent on PSKR signaling.

Fig. S7. ABA accumulation under mannitol stress in roots.

FIG. S8. Differences in ABA concentration were measured using the FRET-based ABA sensor ABAleon.

FIG. S9. Regulation of *NCED* expression by mannitol and ABA.

FIG. S10. Regulation of *UGT71B* and *BG* transcripts by mannitol, ABA and PSKR signaling.

FIG. S11. Regulation of *CYP707A* ABA-8’-hydroxylase transcripts by mannitol, ABA and PSKR signaling.

## ACKNOWLEDGEMENTS

Hagen Stellmach (IPB Halle) is acknowledged for helping in ABA measurements and Prof. Wolfgang Bilger (University of Kiel) for help with PAM imaging. We are grateful to Rainer Waadt and Karin Schumacher (University Heidelberg) for providing the *ABAleon2.1* line and the plasmid *barII-UT-ABAleon2.1*. This work was funded by the Deutsche Forschungsgemeinschaft through grants SA495/8-1 and SA495/8-2.

## AUTHOR CONTRIBUTIONS

MS conceived the project. MS, KR and MDS designed the experiments. BH performed ABA measurements and data analysis. KR and MDS performed experiments and analyzed data. MS wrote the manuscript with contributions from KR, MDS and BH.

## DATA AVAILABILITY STATEMENT

All data supporting the findings of this study are available within the paper and within its supplementary data published online.

