## Supplementary Data for "PSK signaling controls ABA homeostasis and signaling genes and maintains shoot growth under osmotic stress"

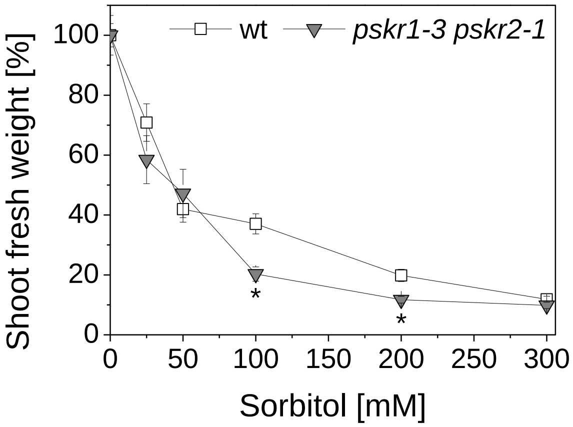


**Supplementary Figure S1. PSKRs confer enhanced resistance to sorbitol.** Four-day-old seedlings were transferred to medium supplemented with different sorbitol concentrations and analyzed after another 3 weeks. Mean ± SE of shoot fresh weight of PSK receptor double knockout mutants on sorbitol are given in percent with values under control conditions set to 100%. Asterisks indicate significant differences between genotypes (Mann-Whitney test, $p<0.05$, n≥18, 3 independent experiments).

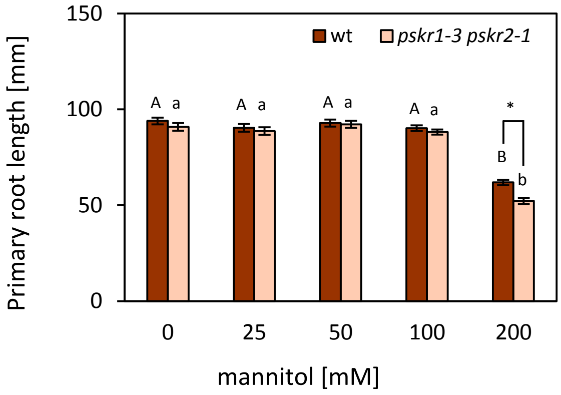


**Supplementary Figure S2. PSK receptor signaling is required to maintain root growth under severe osmotic stress.**

Four-day-old wild type and *pskr1-3 pskr2-1* seedlings were transferred to media containing mannitol as indicated. Primary root lengths of seedlings grown for 7 days after transfer. Significantly different values within a genotype are denoted by different letters (Kruskal-Wallis, Tukey's ‚test, $p<0.05$, $n \geq 36$, 3 independent experiments). Asterisk indicates significant differences between genotypes (Mann-Whitney test, $p<0.05$).

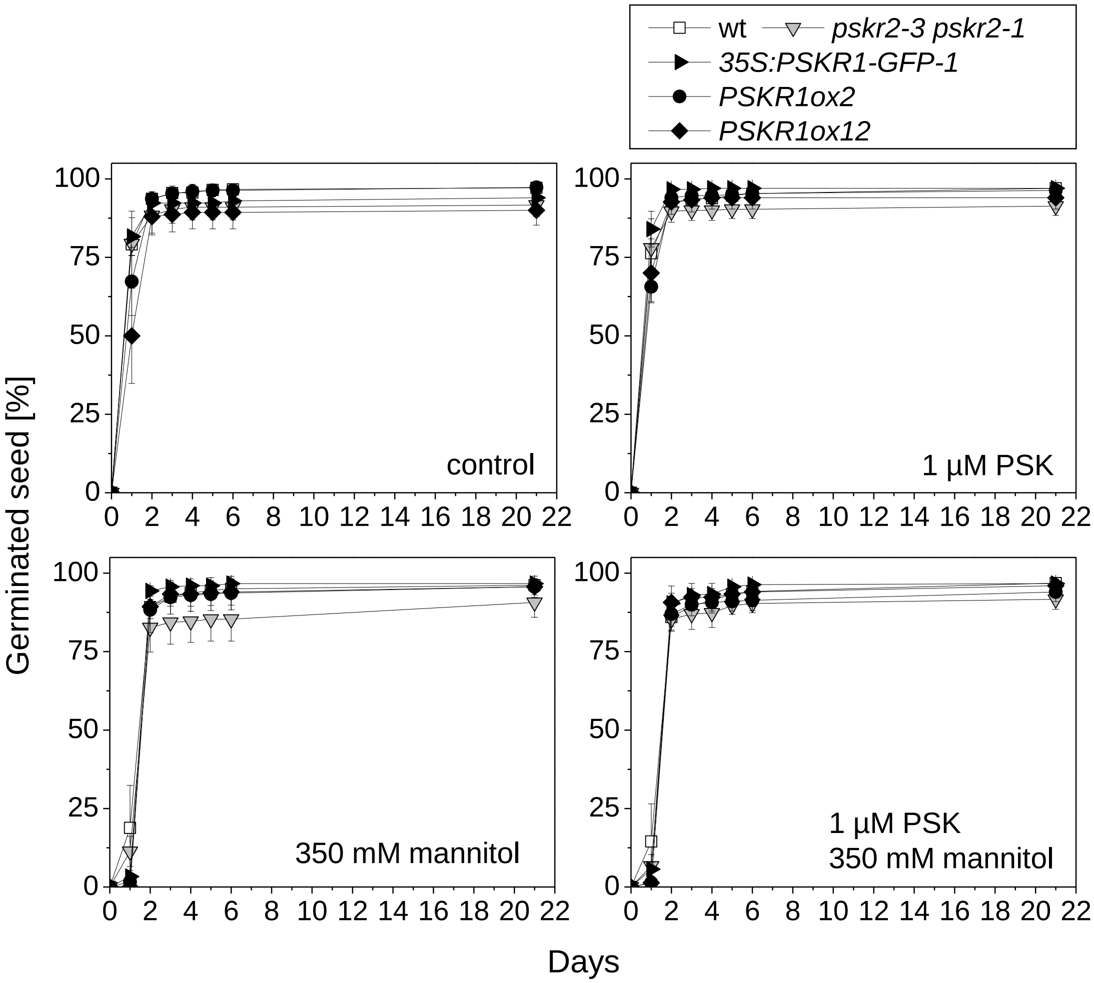


**Supplementary Figure S3. Inhibition of seed germination by mannitol is not dependent on PSK receptor signaling.**

Germination rates were determined in response to 1 µM PSK and 350 mM mannitol. Values are means (± SE) from three independent biological experiments each with 100 seeds analyzed. Seeds with a visible radicle were scored as germinated.

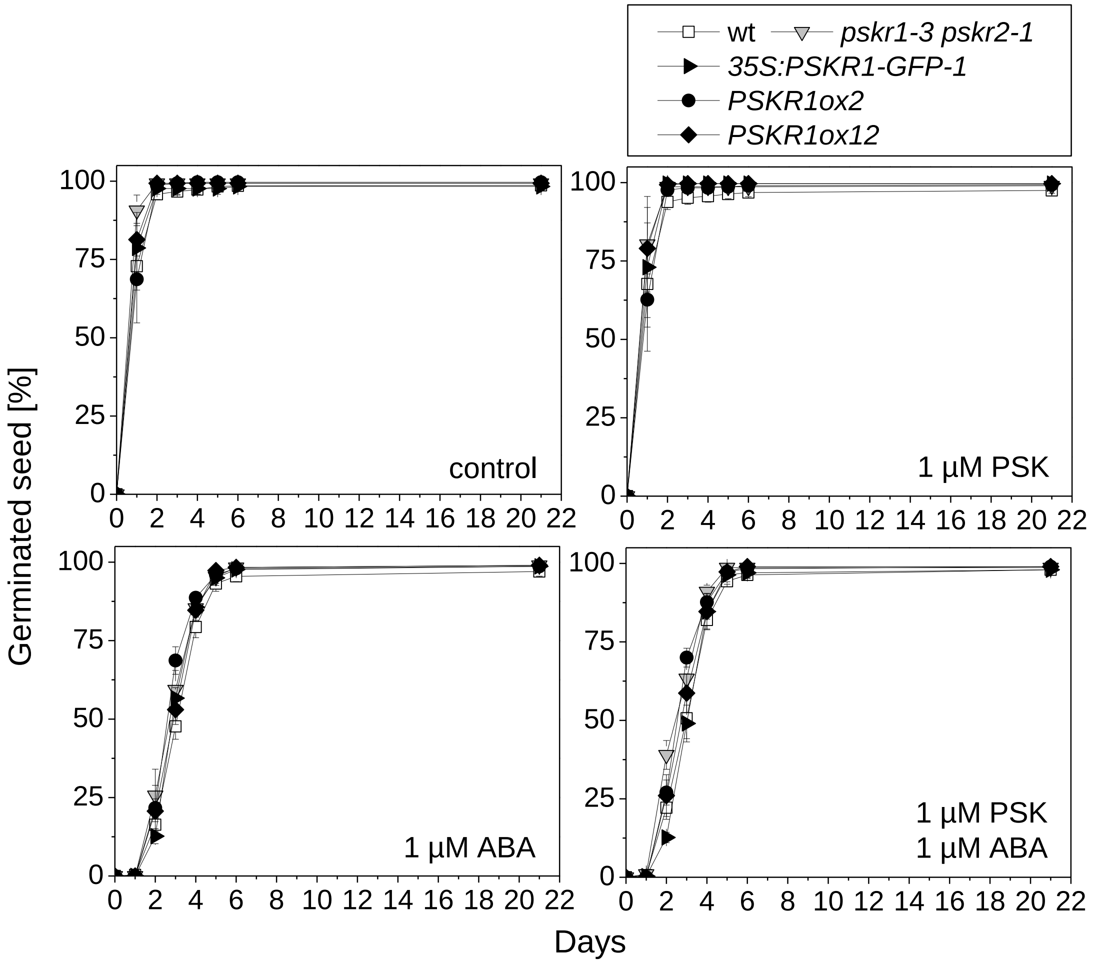


**Supplementary Figure S4. Inhibition of seed germination by ABA is not dependent on PSK receptor signaling.**

Germination rates were determined in response to 1 µM PSK and 1 µM ABA. Values are means (±SE) from three independent biological experiments each with 100 seeds analyzed. Seeds with a visible radicle were scored as germinated.

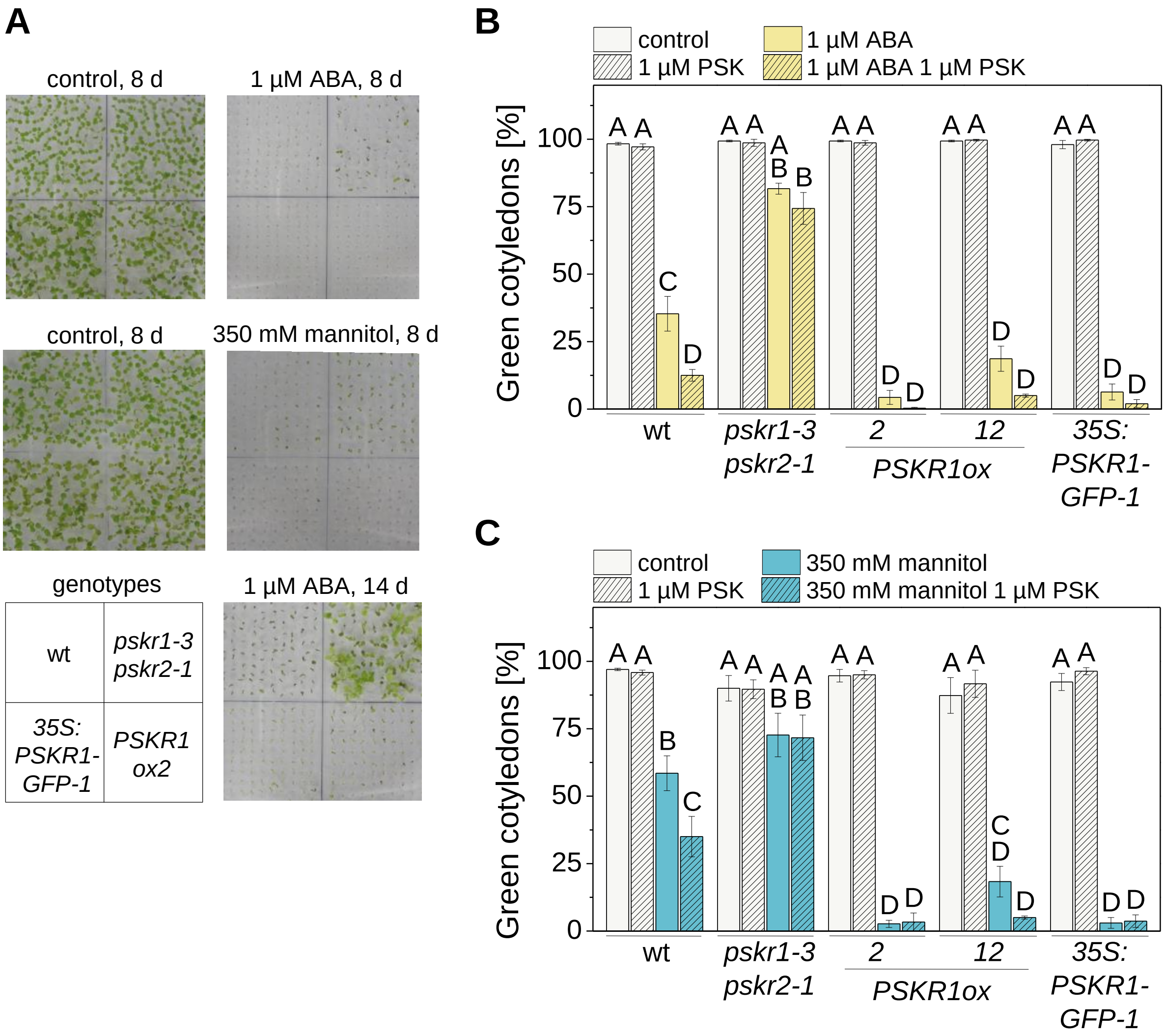


**Supplementary Figure S5. Repression of early seedling development by ABA or mannitol is mediated by PSK receptor signaling.**

**A** Cotyledon greening of wild type, *pskr1-3 pskr2-1*, and *PSKR1* overexpressing seedlings exposed to 1 µM ABA or 350 mM mannitol or no treatment for the times indicated. The scheme on the lower left side indicates the genotypes grown on each plate

**B** Cotyledon greening of 8-day-old wild type seedlings, *PSKR* knockout and of three independent *PSKR1* overexpressing lines treated with 1 µM PSK, 1 µM ABA, or both. Data are means ±SE of 3 independent experiments each evaluating 100 seeds per genotype. Different letters indicate significantly different values (one-way ANOVA, Tukey's test, $p<0.05$).

**C** Cotyledon greening of 8-day-old wild type seedlings, *PSKR* knockout and of three independent *PSKR1* overexpressing lines treated with 1 µM PSK, 350 mM mannitol, or both. Data are means ±SE of 3 independent experiments each evaluating 100 seeds per genotype. Different letters indicate significantly different values (one-way ANOVA, Tukey's test, $p<0.05$).

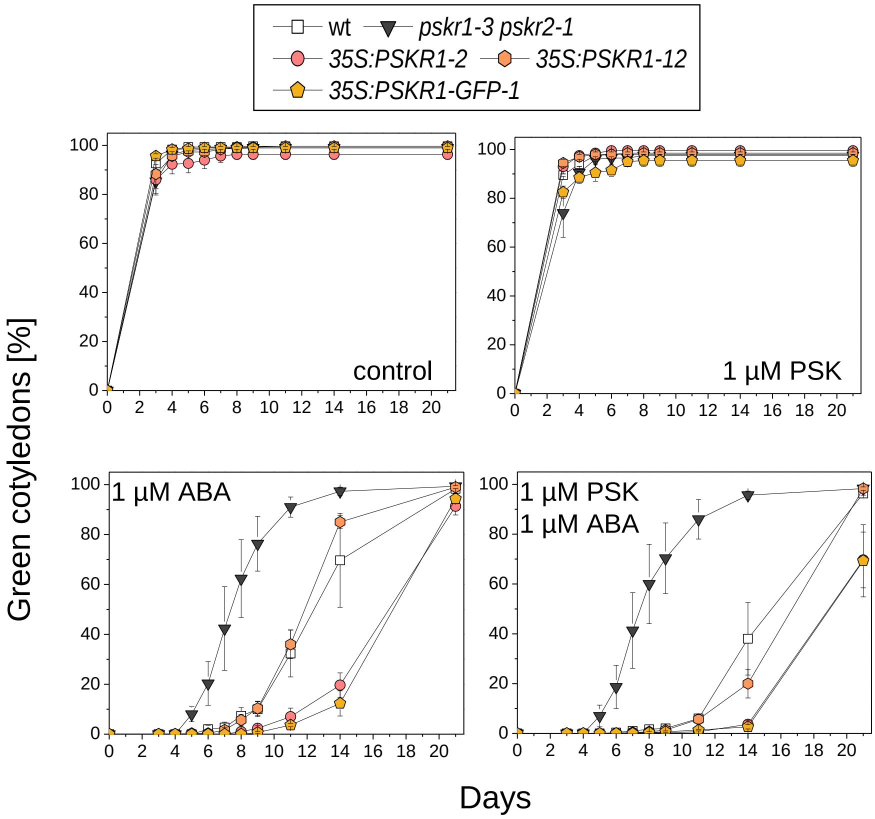


**Supplementary Figure S6: Inhibition of cotyledon greening by ABA is partially dependent on PSKR signaling.**

Time course analysis of cotyledon greening in response to ABA and PSK. Values are means ± SE of three independent biological experiments, each evaluating 100 seedlings. Seedlings with green expanded cotyledons were scored.

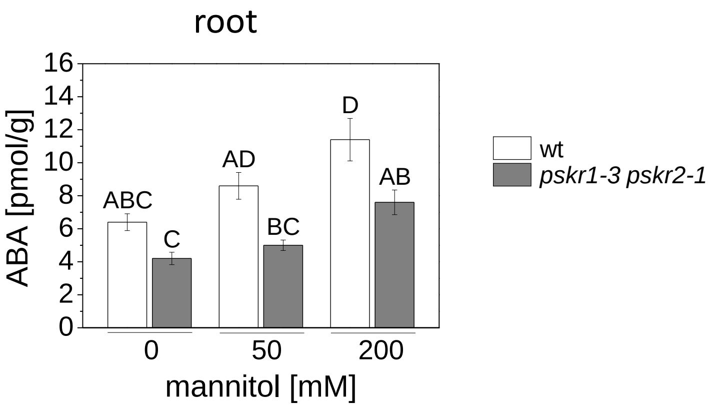


**Supplementary Figure S7. ABA accumulation under mannitol stress in roots.** Four-day-old wild type and *pskr1-3 pskr2-1* seedlings were transferred to 0 mM, 50 mM or 200 mM mannitol and ABA levels in the root were determined after 7 days. Different letters indicate statistically significant differences (Kruskal-Wallis, Tukey‘s test, $p<0.05$, 5 biological replicates).

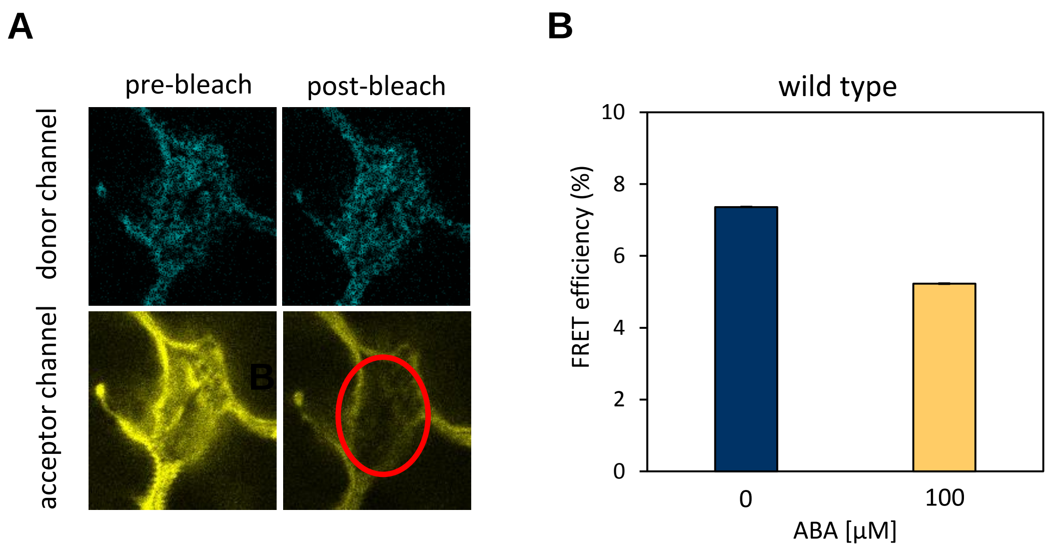


**Supplementary Figure S8. Differences in ABA concentration were measured using the FRET-based ABA sensor ABAleon.**

(**A**) Acceptor photobleaching FRET in wild type Arabidopsis leaves co-expressing the mTurquoise (donor) and cpVenus173 (acceptor) construct. The images were excited sequentially at 458 and 514 nm and emission recorded with adequate filter sets. The region of the cell indicated by a red circle was photobleached at 514 nm for 10 sec. Post-bleach images were captured at 458 nm excitation. FRET is visualized as an increase in mTurquoise fluorescence following cpVenus173 photobleaching, which shows a distinct increase in donor post-bleaching; scale bar = 20 µm. **(B)** Seven-day-old wild type seedlings were transferred to media with or without 100 µM ABA and were analyzed after 1 day. FRET efficiency of the ABA sensor ABAleon was measured using acceptor photobleaching.

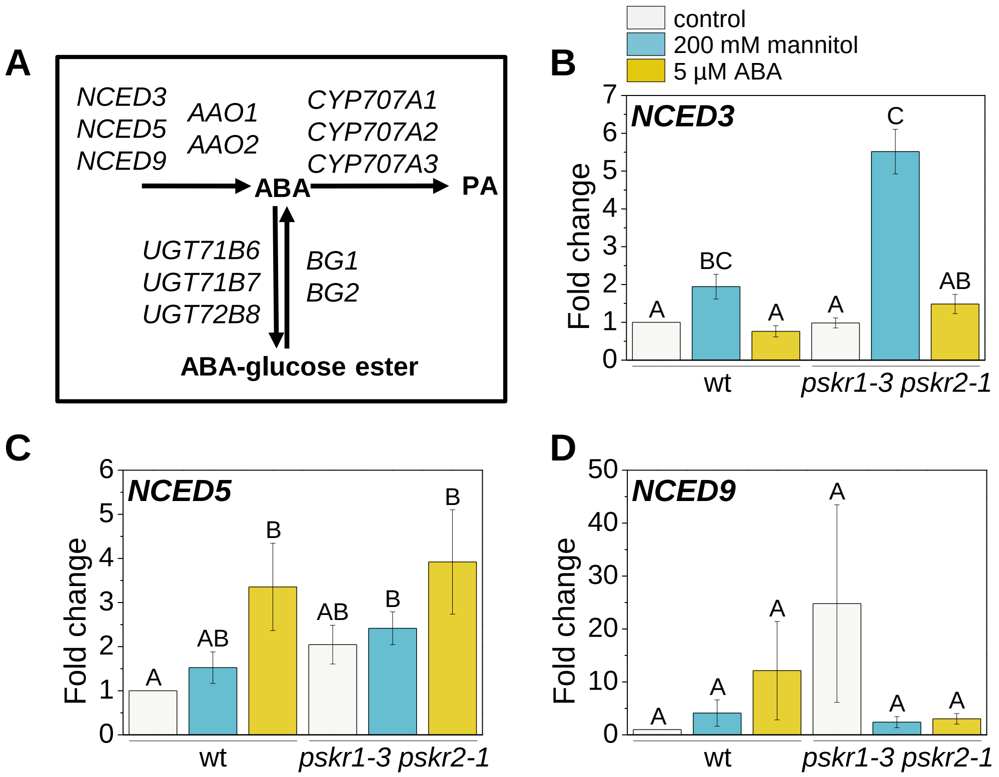


**Supplementary Figure S9. Regulation of *NCED* expression by mannitol and ABA.**

Seven-day-old wild type and *pskr1-3 pskr2-1* seedlings were exposed to 200 mM mannitol or 5 µM ABA for 24 h to analyze relative *NCED3, 5* and *9* transcript levels in true leaves.

**A** Overview of genes encoding for enzymes of ABA synthesis (*NCED*s, AAOs), degradation (*CYP707A*s), inactivation (*UGT71B*s) and mobilization (*BG*s). PA=phaseic acid.

**B-D** Changes in *NCED3*, *NCED5* and *NCED9* transcript levels in response to mannitol, ABA and genotype. Values are means (±SE) with different letters indicating significant differences (Kruskal-Wallis, $p<0.05$; n=6, 3 biological replicates).

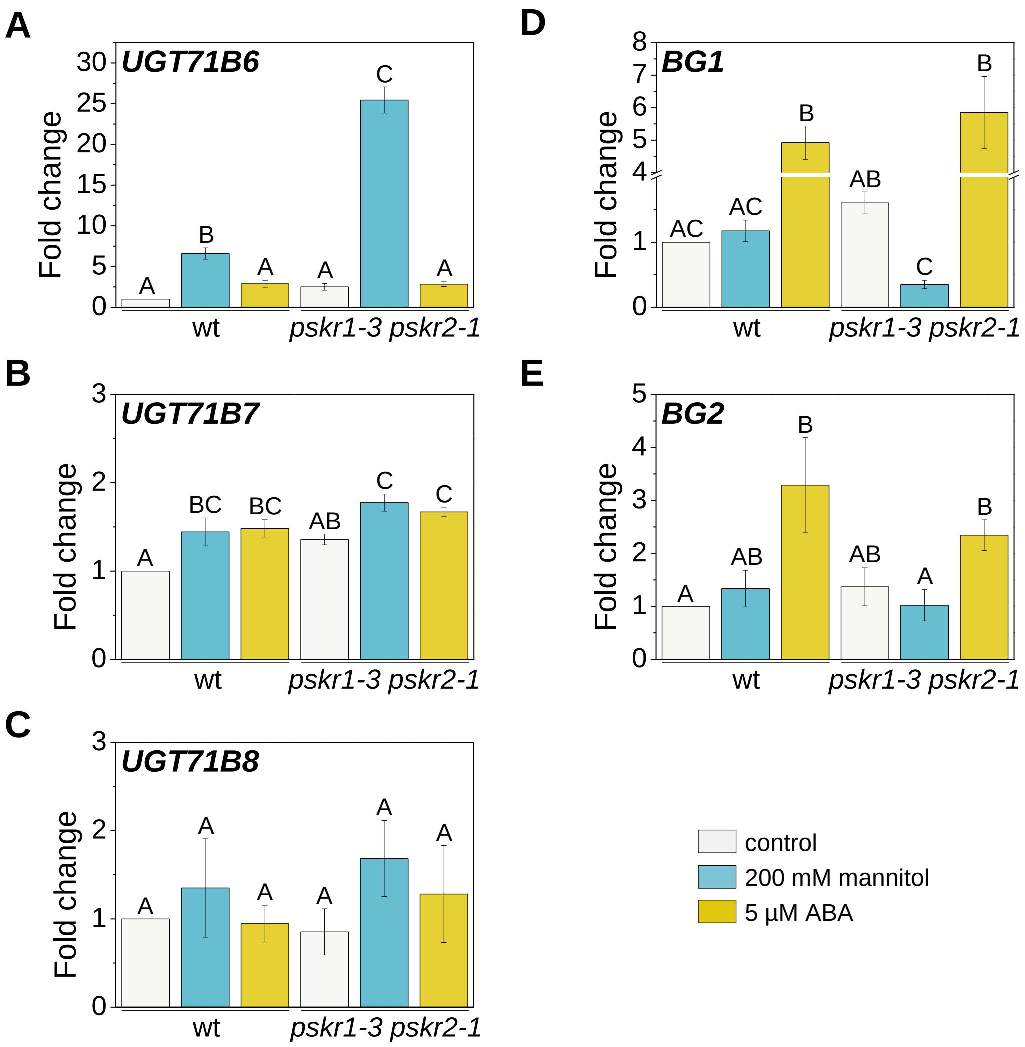


**Supplementary Figure S10. Regulation of *UGT71B* and *BG* transcripts by mannitol, ABA and PSKR signaling.**

Seven-day-old wild type and *pskr1-3 pskr2-1* seedlings were exposed to 200 mM mannitol or 5 µM ABA for 24 h for RT-qPCR analysis in true leaves. Values are means (± SE) of 3 biological replicates with different letters indicating significant differences

**A-C** Relative mRNA levels of *UGT71B6-8* in response to genotype and treatment (Kruskal-Wallis, Tukey‘s test, $p<0.05$).

**D, E** Relative mRNA levels of *BG1 and 2* in response to genotype and treatment (one-way ANOVA, Tukey‘s test, $p<0.05$).

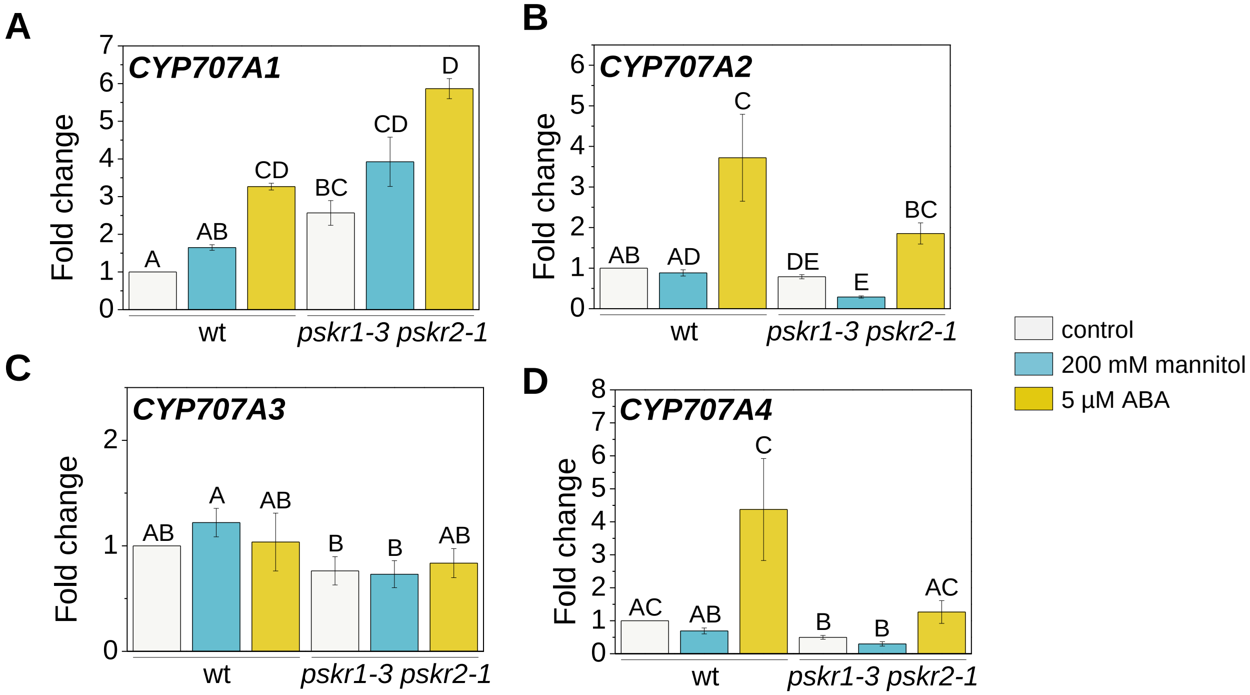


**Supplementary Figure S11. Regulation of *CYP707A* ABA-8'-hydroxylase transcripts by mannitol, ABA and PSKR signaling.**

Seven-day-old wild type and *pskr1-3 pskr2-1* seedlings were exposed to 200 mM mannitol or 5 µM ABA for 1 day and analyze by RT-qPCR for gene expression in true leaves.

**A-D** Relative *CYP707A1-4* transcript levels in response to genotype and treatment (Kruskal-Wallis, Tukey's test, $p<0.05$, n=6, 3 biological replicates). Values are means (±SE) with different letters indicating significant differences (Kruskal-Wallis, Tukey's test, $p<0.05$, n=6; 3 biological replicates).

**Supplementary Table S1.** Primer sequences used for qRT-PCR. for: forward, rev comp: reverse complement.

| Gene name | Accession number | Orientation | Sequence 5' - 3' |
|---|---|---|---|
| *ACT2* | At3g18780 | for | ACATTCCAGCAGATGTGGATCTC |
| | | rev comp | GATCCCATTCATAAAACCCCAGC |
| *GAPC1* | At3g04120 | for | GATTCTACAATGGCTGACAAGAAGA |
| | | rev comp | ATGAAGGGGTCGTTGACAGC |
| *PSKR1* | At2g02220 | for | AGCGAGGTTTTCGATCCGTT |
| | | rev comp | CTGTTGAGTCGTTGGCCTCT |
| *PSKR2* | At5g53890 | for | GTTCGCTCAAGAGGGAGATCA |
| | | rev comp | CCGGTGCCCGTGTTTGTAT |
| *PSK1* | At1g13590 | for | CCATCTCCCACCACACATGA |
| | | rev comp | TCGGTGTGAGCAGCTACTGT |
| *PSK2* | At2g22860 | for | ACTGCTGCCGATCCATGTAA |
| | | rev comp | GCGACCAAAGTCCTCCTCAT |
| *PSK3* | At3g44735 | for | CTTGTGCCTGGCAGTTCTCT |
| | | rev comp | ACTGAGTCCTCTTCCACTGATG |
| *PSK4* | At3g49780 | for | GACAACCACCGCTTTGTCCA |
| | | rev comp | GGTGTGAAGAACAAGGCTTCG |
| *PSK5* | At5g65870 | for | CTCTGCTCCACGCTAACACA |
| | | rev comp | TCTTCTCCAACACCTTCACAGA |
| *CYP707A1* | At4g19230 | for | GAACCCACTCGTGTCCTGG |
| | | rev comp | GTTTTGGGGAAGCGCGAATG |
| *CYP707A2* | At2g29090 | for | GGATCAAAAACGCAACGGCT |
| | | rev comp | GCTCCCTGGAGACTTCTTGG |
| *CYP707A3* | At5g45340 | for | CCGTAGCTCCTCCACGAAAC |
| | | rev comp | GGGCTCGAGATCATCACACAT |
| *CYP707A4* | At3g19270 | for | CGAAGCCGAATACATTCATGCC |
| | | rev comp | TCCCTTCACTTCCCATCGGAAA |
| *AAO1* | At5g20960 | for | GCAAACAGGAGGACCTGTGA |
| | | rev comp | TTACTTCAACCTCGCTCGC |
| *AAO2* | At3g43600 | for | CTGCGGTGAAGGAGGATGT |
| | | rev comp | ACGCTGCAGAGGAGTGTAAG |
| *ABI1* | At4g26080 | for | CATGTCGAGATCCATTGGCGA |
| | | rev comp | TCACACGCTTCTTCATCCGT |
| *ABI2* | At5g57050 | for | AGATCCATTGGCGATAGATACCTT |
| | | rev comp | CCAAATCGGCACACTTCTTCGT |
| *BG1* | At1g52400 | for | TGCCGAGAATGGATACGGAG |
| | | rev comp | TTGTCCTTGCAAATGGCGTC |
| *BG2* | At2g32860 | for | ACTGTAAAAAGGTCGAGGGTGG |
| | | rev comp | TGGTACTCGAGGGATCGTCT |
| *EGM1* | At1g11300 | for | TGTCATGAAGAAAAGAGCGAAAA |
| | | rev comp | GTTGCTGCTGCTAGCACTT |
